# *Expression of* Microcystis *biosynthetic gene clusters in natural populations suggests temporally dynamic synthesis of novel and known secondary metabolites in western Lake Erie*

**DOI:** 10.1101/2022.10.12.511943

**Authors:** Colleen E. Yancey, Fengan Yu, Ashootosh Tripathi, David H. Sherman, Gregory J. Dick

## Abstract

**Summary:** *Microcystis* spp. produces diverse secondary metabolites within freshwater cyanoHABs around the world. In addition to the biosynthetic gene clusters (BGCs) encoding known compounds, *Microcystis* genomes harbor numerous BGCs of unknown function, indicating its poorly understood chemical repertoire. While recent studies show that *Microcystis* produces several metabolites in the lab and field, little work has focused on analyzing the abundance and expression of its broader suite of BGCs during cyanoHAB events. Here, we use metagenomic and metatranscriptomic approaches to track the relative abundance of *Microcystis* BGCs and their transcripts throughout the 2014 western Lake Erie cyanoHAB. Results indicate the presence of several transcriptionally active BGCs that are predicted to synthesize both known and novel secondary metabolites. The abundance and expression of these BGCs shifted throughout the bloom, with transcript abundance levels correlating with temperature, nitrate and phosphorus concentrations, and the abundance of co-occurring predatory and competitive eukaryotic microorganisms, suggesting the importance of both abiotic and biotic controls in regulating expression. This work highlights the need for understanding the chemical ecology and potential risks to human and environmental health posed by secondary metabolites that are produced but unmonitored, as well as the potential discovery of pharmaceutical compounds from cyanoHAB-derived BGCs.

**Originality-Statement of Significance:** *Microcystis spp*. dominate cyanobacterial harmful algal blooms (cyanoHABs) worldwide and pose significant threats to water quality through the production of numerous secondary metabolites, many of which are toxic. While the toxicity and biochemistry of microcystins and several other compounds have been well studied, the broader suite of secondary metabolites produced by *Microcystis* remains poorly understood, leaving gaps in our understanding of their impacts on ecology, human and ecosystem health, or potential pharmaceutical application. In this study, we use metagenomic and transcriptomic datasets to examine the diversity of genes encoding synthesis of secondary metabolites in natural *Microcystis* populations and assess their patterns of transcription in the context of biotic and abiotic conditions in western Lake Erie cyanoHABs. Our results reveal the presence of a large diversity of both known gene clusters that encode toxic secondary metabolites as well as novel ones that encode cryptic compounds. This research highlights the need for targeted studies of the secondary metabolite diversity in western Lake Erie, a vital freshwater source to the United States and Canada.

## Introduction

CyanoHABs, or dense proliferations of cyanobacteria that can discolor water and produce toxins, occur annually and globally, and are expected to increase in severity with climate change (Huisman *et al*., 2018; Ho *et al*., 2019; Griffith and Gobler, 2020). These blooms, which can persist through the early summer into late fall in temperate environments, produce a range of secondary metabolites that can be deleterious to ecosystem function and human health (Harke *et al*., 2016; Watson *et al*., 2016; Steffen *et al*., 2017). CyanoHABs result largely from anthropogenic eutrophication via nutrient runoff (Heisler *et al*., 2008; Huisman *et al*., 2018; Paerl *et al*., 2018) and are expected to become more toxic with increasing nutrient loading and continued, intensifying climate change (Chapra *et al*., 2017; Huisman *et al*., 2018; Krausfeldt *et al*., 2019).

*Microcystis* is a non-N-fixing, potentially toxic cyanobacterium that often dominates freshwater cyanoHABs on every continent except Antarctica (Harke *et al*., 2016). Blooms made up of largely *Microcystis* are expected to expand and increase in severity over the next several years (Watson *et al*., 2016; Huisman *et al*., 2018). This expansion is a concern because *Microcystis* produces diverse bioactive secondary metabolites with an array of ecological and physiological functionalities (Welker and von Döhren, 2006a; Kehr *et al*., 2011; le Manach *et al*., 2019). Many *Microcystis* secondary metabolites possess toxic or inhibitory properties toward diverse cell types (Wiegand and Pflugmacher, 2005; Welker and von Döhren, 2006a; Pearson *et al*., 2016) and/or antibiotic, antifungal, or cytotoxic properties that may be useful in pharmaceutical discovery (Ehrenreich *et al*., 2005; Dittmann *et al*., 2015; Newman and Cragg, 2020). Secondary metabolites likely provide *Microcystis* with ecological advantages and may influence cyanoHAB community composition (Ehrenreich et al., 2005; Dittmann et al., 2015; Pérez-Carrascal et al., 2019; Dick et al., 2021). Despite their significance to ecology, human, and ecosystem health, the physiological and environmental controls on biosynthesis of *Microcystis* secondary metabolites remain poorly understood.

Microcystins, the most extensively studied family of secondary metabolites produced by *Microcystis* (Hughes *et al*., 1958; Carmichael, 1992; Tillett *et al*., 2000), are structurally related hepatotoxins that are responsible for several drinking crises (Kotut *et al*., 2006; Qin *et al*., 2010; Steffen *et al*., 2017) and livestock poisoning events (Stewart *et al*., 2008) around the world. However, *Microcystis* genomes also contain BGCs (Dittmann *et al*., 2015; Pérez-Carrascal *et al*., 2019; Pearson *et al*., 2020) encoding additional secondary metabolites such as aeruginosins, anabaenopeptins, cyanobactins, cyanopeptolins, microginins, and microviridins (Welker and von Döhren, 2006a; Kehr *et al*., 2011), which can be toxic and may occur in the environment and drinking water treatment plants at frequencies and concentrations that equal or exceed those of microcystins (Beversdorf *et al*., 2017; Janssen, 2019; Kust *et al*., 2020). In addition, many BGCs identified from *Microcystis* genomes do not have associated products, and thus have been designated “orphan” clusters (Humbert *et al*., 2013; le Manach *et al*., 2019), highlighting the need to identify and determine the structural characteristics and biological properties of these unknown compounds. While *Microcystis* genomes harbor a diverse and variable suite of BGCs (Pérez-Carrascal et al., 2019; Dick et al., 2021), and some metabolites have been shown to be expressed and produced in culture (Dittmann *et al*., 2015), little is known about the occurrence or expression of these *Microcystis* BGCs in natural environments. Understanding their distribution and expression in relation to the biotic and abiotic environment may shed light on the ecological role of these molecules and inform risks to public health.

In this study we used metagenomics and metatranscriptomics to assess the abundance, diversity, and expression of BGCs in *Microcystis* populations in a time series at three stations during the 2014 western Lake Erie bloom, which show a succession of *Microcystis* strains alongside changing nutrient availability, microcystin concentrations, and microbial communities (Berry *et* a*l*., 2017; Yancey *et al*., 2022). Western Lake Erie is subject to annual cyanoHABs that have intensified in recent decades (Steffen *et al*., 2014; Watson *et al*., 2016), and the 2014 bloom caused a drinking water crisis in which Toledo residents lost access to potable water due to dangerously high levels of microcystins (Steffen *et al*., 2017). While microcystins are heavily monitored and studied in these waters, monitoring of “other” secondary metabolites has been limited to saxitoxin, anatoxin, and cylindrospermopsin (Lake Erie Programs | Ohio Environmental Protection Agency). Motivated by the known diversity of *Microcystis* BGCs and the influence of nutrient stoichiometry on biosynthesis of secondary metabolites (van de Waal *et al*., 2009; Wagner *et al*., 2019), we hypothesized that the relative abundance and transcription of BGCs would vary across the bloom season as nutrient availability changes.

## Experimental Procedure

### Study Site and Sample Collection

Samples were collected weekly from various NOAA Great Lakes Environmental Research Laboratory (GLERL) sampling stations throughout the western Basin of Lake Erie from mid-June through late October 2014 (Cooperative Institute for Great Lakes Research). The long-term NOAA GLERL stations selected for sampling were WE2, WE4, and WE12. WE2 is close to the inlet for the Maumee River inlet (41° 45.743’N, 83° 19.874’ W), WE4 is considered an offshore site closer to the center of the basin (41° 49.595’N, 83° 11.698’W), and WE12 is proximal to the Toledo drinking water inlet (41° 42.535’N, 83° 14.989’W) (Fig. 1A).

**Figure 1:**
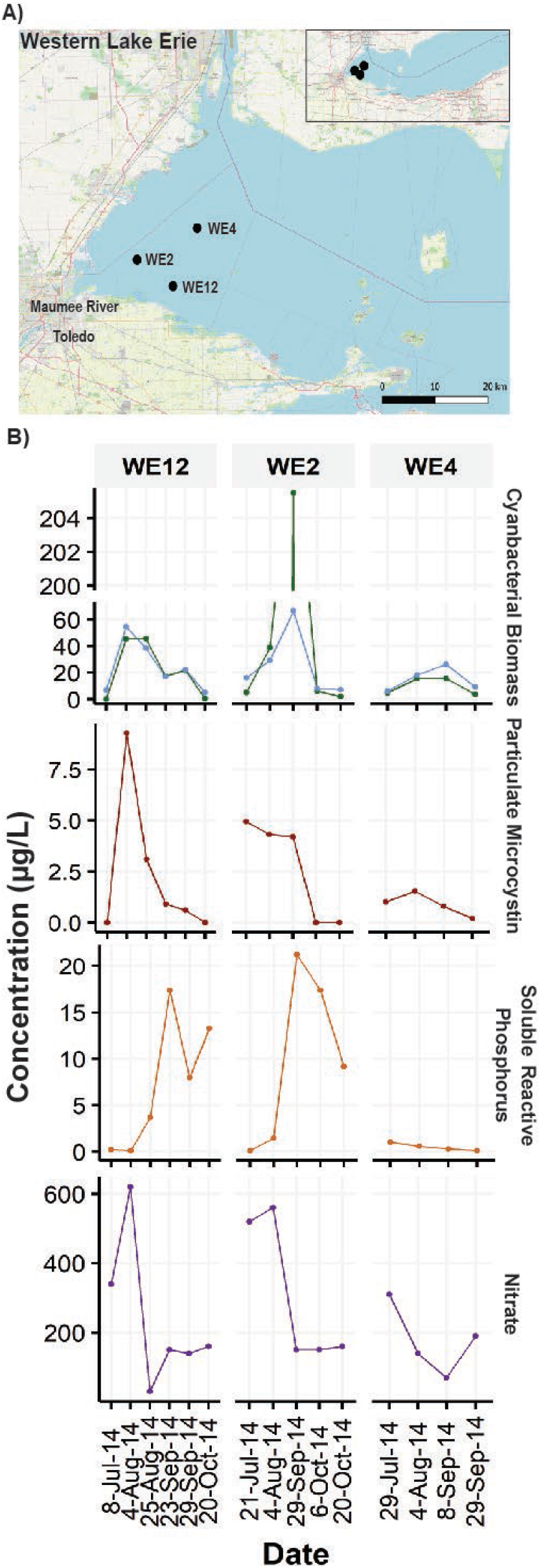
Overview of the 2014 W. Lake Erie cyanoHAB. A) Map of western Lake Erie and the sampling sites used for this study. WE2 and WE12 are close to the Ohio coast and Maumee River and are considered nearshore stations, while WE4 is located more centrally in the basin and considered offshore. B) Various bloom metrics for the 2014 bloom including cyanobacterial biomass as measure by both chlorophyll α (blue) and phyocyanin (green) concentrations, as well as measured concentrations of particulate microcystins, soluble reactive phosphorus, and nitrate. Figure 1A was generated via QGIS using the Open Street Map (OSM) as a basemap (https://wiki.osmfoundation.org/wiki/Main_Page).

Twenty liters of depth integrated samples were collected from each station for biological and chemical analysis. Integrated depths were defined as the surface of the water to 1m above the lake floor. While samples were collected, measurements of pH and water temperature were performed as well. To capture *Microcystis* aggregates, 2L of integrated depth collected water were filtered through a 100 µm polycarbonate mesh filter. The biomass on the filter was then collected and filtered through a 0.22 µm filter. The biomass on the 0.22 µm filter was preserved in 1 mL of RNALater™ (Invitrogen™, Ambion™) and placed on ice. Upon arrival to the lab, samples were stored in the -80°C freezer until DNA and RNA extractions could be completed.

### Sample Processing and Sequencing

Processing and sequencing of these samples was described previously (Yancey *et al*., 2022). Briefly, DNA was extracted with the Qiagen DNeasy Blood and Tissue Extraction Kit (Qiagen, Hilden, Germany) and quantified with the Quant-iT™ PicoGreen™ dsDNA Assay Kit (Eugene, OR, USA). The Qiagen RNeasy Kit (Qiagen, Hilden, Germany) was used to complete RNA extraction. Shotgun DNA and RNA sequencing was done at the University of Michigan DNA Sequencing Core on the Illumina® HiSeq™ platform (2000 PE 100, Illumina, Inc., San Diego, CA, USA).

### Bioinformatic Analysis

Shotgun reads were assembled *de novo* to recover *Microcystis* Metagenome Assembled Genomes (MAGs). Metagenomic raw read sequences obtained from the University of Michigan Omics Core were analyzed and processed through the following metagenomic workflow. Raw reads for metagenomic analysis can be found at NCBI BioProject Accession: PRJNA464361.

The IMG-JGI supported software, BBTools was used to complete trimming, adaptor removal, and quality check of raw reads. Quality checked (QC) reads were normalized to 100x coverage using bbnorm from the BBTools package (Bushnell). 100x coverage was chosen based on the assembly quality of the following housekeeping genes: a global nitrogen regulator *ntcA*, the beta subunit phycocyanin gene *cpcB*, the photosystem I P700 chlorophyll apoprotein *psaB*, the photosystem II P680 reaction center D1 protein *psbA2*, as well as the assembly quality of the *mcy* operon. Assembly quality was assessed based on contiguous sequence length, and De Bruijn Graph assembly visualized in Bandage (Wick *et al*., 2015).

*De novo* assemblies were generated for each sample using MEGAHIT (Li *et al*., 2015). To obtain differential coverage for binning software, mapping of QC reads was completed using Bowtie2 (Langmead and Salzberg, 2012). A multiple binning approach was used in which bins generated from Concoct, Metabat, Tetra ESOM, and Vizbin were assessed and chosen with DASTool (Ultsch and Mörchen, 2009; Alneberg *et al*., 2013; Kang *et al*., 2015; Laczny *et al*., 2015; Sieber *et al*., 2018). To improve accuracy of binning software, differential coverage of all 15 samples was used by mapping reads from each sample to each assembly using Bowtie2. The minimum cut off for contiguous sequence length was 1 kb for Concoct, 1.5 kb for Metabat, and 3.5 kb for Tetra ESOM and Vizbin. The cut off for Metabat was selected based on minimum cut off length the program required and that for ESOM and Vizbin was selected based on clarity of maps. *Microcystis* MAGs were assessed and manually refined using Anvi’o v. 5, “Margaret” (Eren *et al*., 2015). Completion and strain heterogeneity of each MAG was assessed using CheckM with default parameters (Parks *et al*., 2015) as well as metrics generated through Anvi’o outputs. MAGs were mined for BGCs with antiSMASH 6.0 using a full Hidden Markov Modelled search, orthologous gene prediction, gene border prediction, PFAM analysis, and COG identification (Blin *et al*., 2021). For further BGC annotation, identified BGC sequences from all *Microcystis* MAGs found by antiSMASH were run through PRISM (Skinnider *et al*., 2020) software as well.

For downstream analysis, BGC sequences identified by antiSMASH from all *Microcystis* MAGs were pooled together and deduplicated using the dedupe tool of the BBTools package (Bushnell). Dedupe was set to deduplicate sequences that were at least 90% identical and had no more than 10 substitutions or deletions. This deduplicated set of BGC clusters was used for subsequent analysis with special attention to specific clusters discussed in greater detail. BGC sequences longer than 10,000 bp were saved for future gene and expression quantification calculations. In addition to these BGCs we also closely examined three identified cyanobactin BGC clusters that were ∼7 kb in length. To compare the similarities between clusters of the same class or identification, the clinker software (Gilchrist and Chooi, 2021) was used to generate gene maps to visually inspect for similarities and differences in gene cluster structure.

### BGC Cluster Quantification

To estimate the relative abundance of BGC clusters the following steps were completed. QC metagenomic reads were aligned to all identified BGC clusters found in any cyanobacterial MAG (including *Microcystis, Anabaena, Pseudanabaena*, and *Cyanobium)* generated from this sample set using Basic Local Alignment Search Tool (BLAST) v. 2.8.1 (Altschul *et al*., 1990).Using all identified BGC clusters expanded our database of biosynthetic genes to allow for competitive mapping. Reads were kept for quantification if they were 95% identical to the query sequence, and at least 80% in alignment length. If reads identically mapped to more than one sequence, i.e., they had identical bit scores, the read count was divided among the total number of ambiguously mapped regions using the blast_hit_counts.sh script found at https://github.com/Geo-omics/scripts/tree/master/scripts. Reads mapped to a BGC were then normalized by BGC length.

To compare across samples, reads were also normalized by the 16S rRNA gene V4 variable region. This is based on the assumption that all *Microcystis* cells contain at least one 16S rRNA gene, but do not contain all BGCs; therefore, we used this ratio to estimate the relative abundance of BGCs in *Microcystis* populations. All V4 regions for the entire Phylum of Cyanobacteria were used in the mapping databases to ensure competitive, sensitive, and accurate mapping. These sequences were pulled from the SILVA v.1381.1 database (Quast *et al*., 2013) accessed in February 2021. This database can be found at: https://github.com/ceyancey/mcyGenotypes-databases. QC metagenomic reads were aligned to this database using BLAST and kept for downstream analysis if they met the same parameters listed above. These alignment parameters, databases, and mapping tool quality were tested in full to ensure specificity and sensitivity. Reads were then quantified in the same fashion described above and normalized by 16S rRNA V4 region length. The equation below summarizes how relative BGC cluster abundance was calculated:

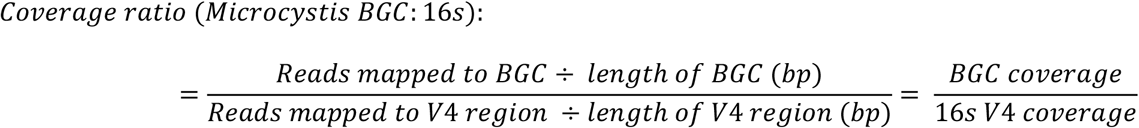

### Biosynthesis Gene Protein Phylogenies

To explore the novelty and relatedness of coarsely annotated PKS clusters from our *Microcystis* MAGs to previously identified biosynthesis pathways, protein phylogenies were generated. BGCs of three polyketide synthase (PKS) types, i.e., T1PKS 1, T1PKS 3, and T3PKS, were chosen for this analysis. For each PKS type, one to two genes that encode for proteins involved in transport, additional synthesis processes, or core biosynthesis (target sequences) were selected. To find related protein sequences, these sequences were searched against the nonredundant protein database on NCBI (accessed July 2022) using BLASTP (Madden, 2013). Protein sequences were used to build phylogenetic trees if they had at least a 70% alignment length and at least 30% identity to the target sequences. Protein sequences were aligned using MUSCLE (Edgar, 2004) and FastTree (Price *et al*., 2009) was used to construct phylogenetic trees. Protein sequences that are ∼30% identity to the target protein sequences were used to root trees. Bootstrap values equal to or greater than 0.5 were reported. PaperBLAST (https://papers.genomics.lbl.gov/cgi-bin/litSearch.cgi) was used to query published manuscripts to investigate whether these protein sequences had been investigated previously.

### BGC Cluster Transcript Abundance

The relative transcript abundance of BGCs *in situ* was also determined.

Metatranscriptomic raw reads can be accessed at NCBI under BioProject Accession: PRJNA370007. Alignments were completed using BLAST with identical parameters mentioned previously. However, to normalize gene expression, reads were mapped to reference genome *Microcystis aeruginosa* PCC 7806SL (Accession: CP020771.1, GI: 1181755937) instead of the V4 region. Competitive genome mapping was achieved by creating a database that contains reference genomes for other common cyanobacteria found in western Lake Erie cyanoHABs: *Anabaena sp*. 90 (GCA_000312705.1), *Cyanobium sp*. NIES-981 (GCA_900088535.1), and *Pseudanabaena sp*. PCC 7367 (GCA_000317065.1). The following equation summarizes calculations completed to relatively quantify BGC expression:

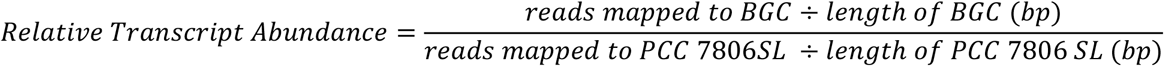

### BGC Expression Correlation Analysis

To assess the relationships between expression and 1) abiotic environmental conditions and 2) abundance of known competitors and predators of *Microcystis*, simple linear regressions were generated. Correlations were generated for all abiotic variables, and for all organisms identified in each metagenome that had at least 200 genes assigned to their taxonomic identification. Organism abundance and analysis was completed at the genus level. Pearson’s correlation coefficient, R, as well as p-values were determined to assess the strength of correlation and the significance of individual variables, respectively. Abiotic environmental condition data was provided by CIGLR and can be found in Table S4. The relative abundance of organisms was determined by aligning QC reads to the UniRef100 database using DIAMOND (Buchfink *et al*., 2021). Relative abundance of each organism was then determine by calculating RPKM of all genes assigned to the taxa using the AnnotateContigs.pl script found at https://github.com/TealFurnholm/Strain-Level_Metagenome_Analysis/wiki/Step-2.-Gene-Annotation-and-Community-Analysis which summarizes the complete community analysis workflow and outputs.

### Data Visualization

All figures were generated using R and R Studio v. 3.5.1 (Allaire, 2015). Packages used for data visualization included ggplot2 (Wickham, 2011), ggthemes (Arnold, 2021), and ggpubr (Kassambara, 2020). To generate the map in Figure 1, QGIS was used using the Quick Map Services Plugin (QGIS Development Team, 2020). The base map used is an open-source resource, the Open Street Map (OSM, https://wiki.osmfoundation.org/wiki/Main_Page). Clinker (Gilchrist and Chooi, 2021) software was used to generate BGC comparison figures observed in Figure S1. Metagenomic and metatranscriptomic raw reads can be found on NCBI under the BioProject Accessions PRJNA464361 and PRJNA370007, respectively.

## Results

### The 2014 Western Lake Erie cyanoHAB

The 2014 western Lake Erie cyanoHAB was notable due to high levels of microcystins at station WE12, the Toledo Drinking water crib, which led to a drinking water crisis event in Toledo, OH, USA. This study used samples and data collected and described previously (Berry et al., 2017; Smith et al., 2021; Yancey et al., 2022). Briefly, samples were collected from three core stations. WE2 and WE12 are close to the Ohio coast and Maumee River inlet and are considered nearshore stations. WE4 is located more centrally in the basin and is considered offshore (Fig 1A). The 2014 bloom consisted of an initial peak in cyanobacteria biomass (as measured via phycocyanin) and microcystin concentrations at nearshore stations in early August. A secondary peak in cyanobacteria biomass, where microcystin concentrations were low, was observed in late September. At the offshore station, WE4, microcystin and phycocyanin concentrations were considerably lower (Fig 1B). Soluble reactive phosphorus (SRP) and nitrate concentrations (NO_3_^−^) were also variable across stations and sampling times, with marked decreases in nitrate during bloom development at stations WE2 and WE12 (Fig 1B).

### Microcystis MAG Generation

Ten *Microcystis* metagenome-assembled genomes (MAGs) were generated from fifteen metagenomic samples from the 2014 western Lake Erie cyanoHAB (Table 1). Samples that failed to generate a *Microcystis* MAG were likely due to low abundance of the organism as seen in early phases of the bloom (early July), or poor assembly of *Microcystis* contiguous sequences due to high strain heterogeneity (Dick, 2018). Most MAGs had high completion (above 90%) and variable redundancy given the high strain heterogeneity observed (Chen *et al*., 2020), except for the MAG generated from August 4^th^, at Station 12, at 40% completion (Table 1). This MAG was kept for further analysis as this sample was taken directly during the time of the Toledo Drinking Water Crisis. Five fragmented BGCs (<5kb) identified in this MAG were removed from further analysis due to incompletion, while one complete BGC (T1PKS 2) was identified and used for subsequent analysis.

**Table 1:** Summary and Statistics of *Microcystis* MAGs

| Station | Date | Completion (%) | Redundancy (%) | N50 | Size (Mbp) | Number of contigs | GC Content (%) |
| --- | --- | --- | --- | --- | --- | --- | --- |
| WE12 | 4-Aug | 40.29 | 2.16 | 2907 | 0.816 | 335 | 39.67 |
|  | 25-Aug | 89.21 | 35.25 | 2353 | 5.99 | 2767 | 43.17 |
|  | 23-Sep | 100 | 4.32 | 8601 | 5.31 | 903 | 42.79 |
|  | 29-Sep | 93.53 | 10.23 | 2548 | 4.27 | 1888 | 43.4 |
|  | 20-Oct | 96.4 | 12.95 | 4155 | 4.92 | 1461 | 42.83 |
| WE2 | 21-Jul | 97.84 | 4.32 | 5000 | 4.27 | 1102 | 42.9 |
|  | 6-Oct | 97.12 | 10.07 | 5061 | 5.39 | 1404 | 43.28 |
|  | 20-Oct | 99.28 | 5.76 | 5472 | 4.69 | 1108 | 42.8 |
| WE4 | 29-Jul | 92.81 | 20.86 | 1786 | 3.93 | 2234 | 43.53 |
|  | 8-Sep | 94.96 | 9.67 | 2668 | 4.29 | 1797 | 43.49 |

### BGC Diversity and Abundance

Nineteen distinct BGCs, representing multiple classes of secondary metabolites, were identified (Fig. 2). The relative abundance of these BGCs, which was estimated by quantifying metagenomic reads mapped to assembled BGCs and normalized to reads mapped to *Microcystis* 16S rRNA genes, showed differences across stations and sampling dates (Fig 2). The abundance of BGCs producing aeruginosins 1 and 2, anabaenopeptin, cyclophane-like 1, micropeptin 3, and microviridin B 1 and 2 increased during later phases of the bloom, especially at nearshore stations during late September through October. BGCs predicted to encode aeruginosins, cyanobactins, microviridins, and micropeptins each yielded multiple distinct clusters with conserved gene content, but gene rearrangements, insertions, and duplications were observed within these clusters (Fig. S1). Gene order and orientation of BGCs encoding aeruginosin and microviridin B were more conserved (Fig S1A & S1C), while rearrangements, insertions, and deletions were more common in the cyanobactin and micropeptin BGCs (Fig S1B & S1D). Deeper annotations for clusters encoding anabaenopeptin, microviridin B, aeruginosin, and a cyanobactin as well as chemical structures of potential congeners produced can be seen in Fig. S2. Some clusters, such as cyclophane-like 1, were rare at most sampling times and stations, except on 20 October when it was one of the most abundant clusters. The cyanobactin clusters, or clusters like those that encode petallamide (Blin *et al*., 2021), shifted in abundance during the bloom with 2 and 3 being more abundant early, and 1 being the most abundant during late phases (Fig. 2).

**Figure 2:**
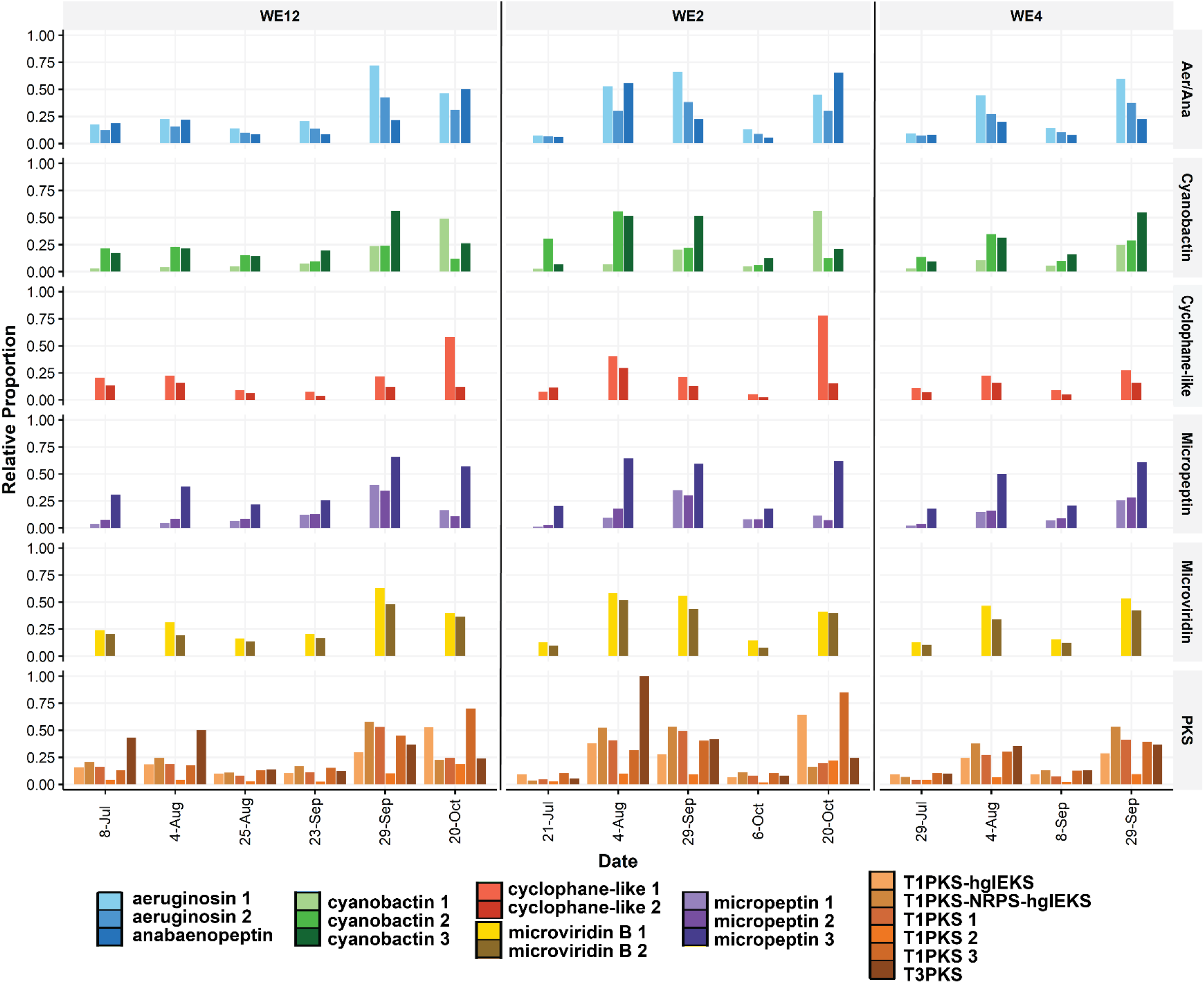
Relative BGC abundance for aeruginosin, anabaenopeptin, cyanobactin, cyclophane-like, micropeptin, microviridin B, and unknown BGC classes. In some cases, relative abundance is above 100% indicating greater copies of BGC genes than 16S rRNA genes

Six cryptic PKS or hybrid-PKS cluster classes were also identified and varied greatly in abundance (Fig. 2). These clusters encode unknown products and have either very low percent similarities (≤40%; Table S1) or no similarity at all to previously described clusters, suggesting high potential for biosynthesis of unique structures. The T3PKS identified reached peak relative abundance at the nearshore stations on 4 August. The T1PKS 3 peaked in abundance at nearshore stations late in the season on 20 October (Fig 2). Four of these cryptic PKS clusters encode biosynthetically interesting enzymes including an enediyne synthase (T1PKS 1) and a tryptophan halogenase (NRPS-T1PKS-hgIEKS) (Fig. 3).

**Figure 3:**
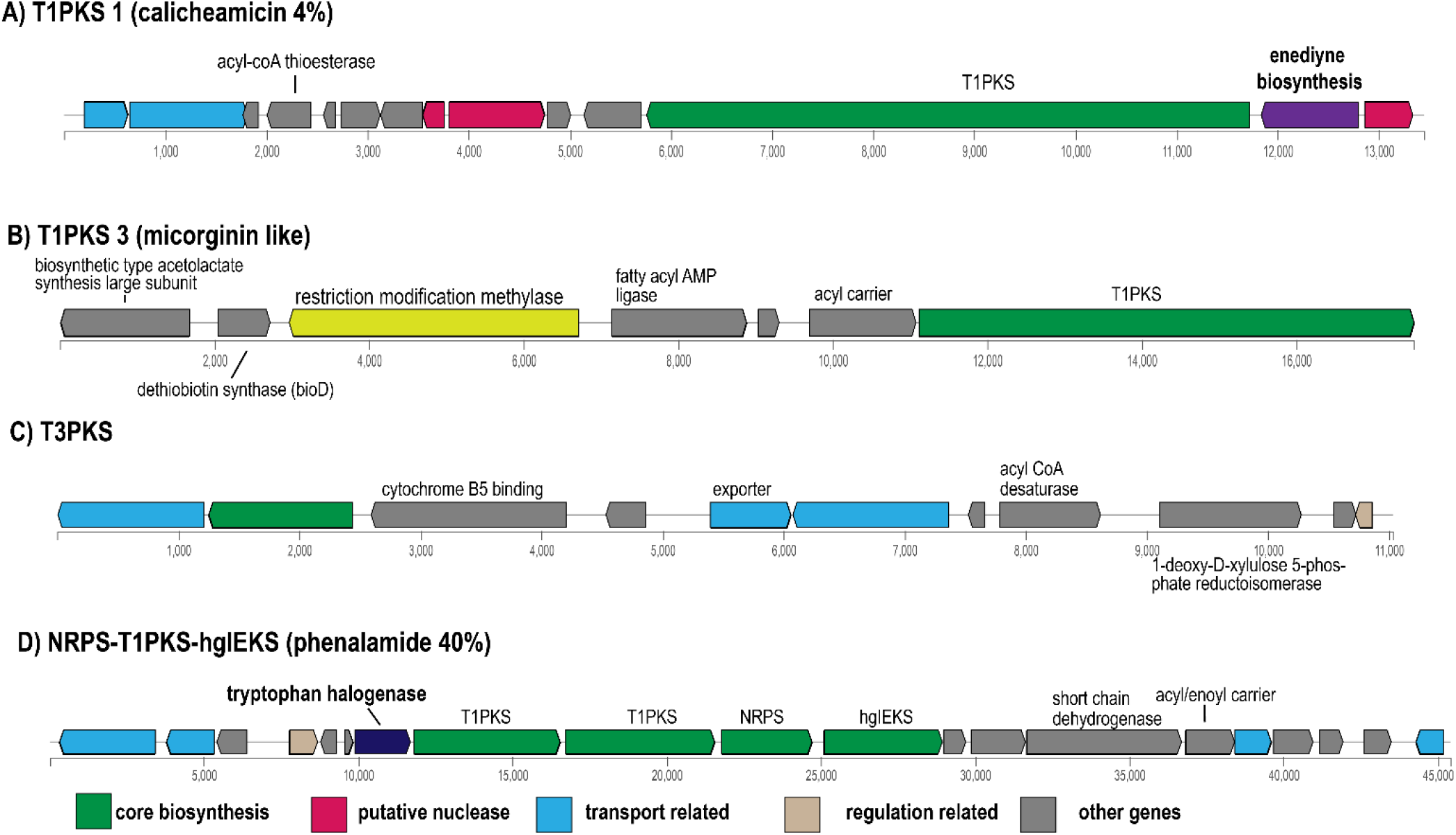
Gene schematics for select, cryptically annotated PKS gene clusters. A) Gene cluster for T1PKS 1 contains transport genes, an enediyne biosynthesis gene, and putative nucleases within the cluster B) T1PKS 3 contains genes that encode for various additional synthesis genes and a regulatory restriction medication methylase. C) The T3PKS cluster contains several transport genes and a cytochrome B5 binding gene. D) The Hybrid NRPS-T1PKS-hgIEKS cluster contains several core biosynthesis and transport genes, and a tryptophan halogenase.

### Protein Phylogenetics for Biosynthesis Genes

Phylogenetic trees of various predicted proteins from within PKS clusters were analyzed for potential insights into biosynthesis pathways. For the T1PKS 3 cluster, the fatty-acyl AMP ligase sequence had high similarity to fatty-acyl AMP ligases from 22 *Microcystis* protein sequences from the nonredundant protein database on NCBI (accessed August 2022), with a bootstrap value of 1 (92.92-97.93% identity, Fig. 4A). To our knowledge, none of these protein sequences have been linked to the synthesis of a known natural product. A sister clade included several sequences encoding secondary metabolite enzymes including fatty-acyl AMP ligases, hybrid NRPS/PKSs, and beta-ketoacyl synthases sequences from a variety of cyanobacteria belonging to the order *Nostacales* (bootstrap = 0.836, Fig. 4A). This clade was highly diverse, with long branch lengths, and had lower percent identity overlap with the Lake Erie MAG sequence (∼59-61%). The core biosynthesis gene identified within the T1PKS 3 cluster was annotated to encode an NRPS/PKS hybrid enzyme, which is closely related to 28 other *Microcystis* NRPS/PKS hybrid enzymes in the NCBI nonredundant protein database (bootstrap = 1, ∼88-99% identity, Fig 4B). Again, none of these *Microcystis* protein sequences are associated with synthesis of a known compound. However, these *Microcystis* sequences clustered within a clade that contained MicA, a type 1 polyketide synthase in the microginin biosynthesis pathway from *Planktothrix prolifica* ((Rounge *et al*., 2009), bootstrap = 1, 81% identity, 95% alignment length, Fig. 4B). The microginin BGC is found in various cyanobacteria including *Planktothrix* spp. (Rounge *et al*., 2009), *Microcystis* spp. (Eusébio *et al*., 2022), and *Anabaena* spp. (Bagchi *et al*., 2016), all of which may be present in western Lake Erie cyanoHABs. Mapping of metagenomic sequence reads to the *Microcystis* microginin BGC (Eusébio *et al*., 2022) showed that the T1PKS 3 shared 93% identity with the BGC described in Eusébio et al. and that the entire BGC is present in western Lake Erie, indicating that the partial contigs is likely due to poor assembly (Fig. S4).

**Figure 4:**
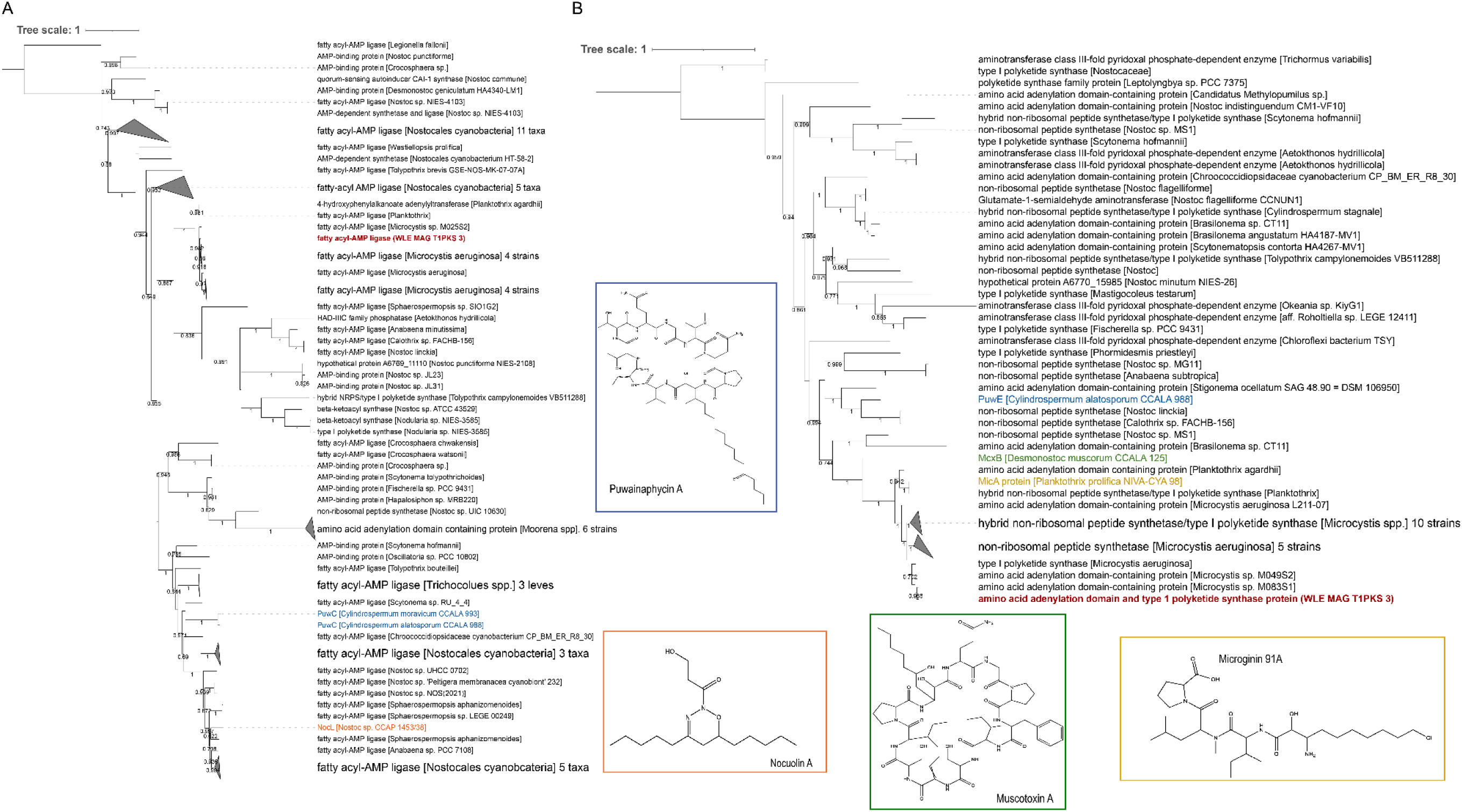
Protein phylogenies for A) a fatty acyl-AMP ligase and B) core biosynthesis NRPS/PKS like synthase from T1PKS 1. The fatty-acyl AMP ligase shares some homology with cyanobacterial synthesis proteins PuwC involved in puwainaphycin synthesis and NocL, involved in nocuolin synthesis. The core biosynthesis protein product from this cluster has protein similarity to genes involved in puwainaphycin (PuwE), muscotoxin (McxB), and microginin (MicA) synthesis.

The enediyne biosynthesis protein sequence from T1PKS 1 was investigated more deeply as enediyne synthesis is a critical step in the formation of anticancer compounds such as calicheamicin (Nicolaou and Dai, 1991). While there were no hits to protein sequences from known biosynthesis pathways, the enediyne protein sequence identified in T1PKS 1 had the highest similarity to 30 *Microcystis* protein sequences in the NCBI nonredundant database, suggesting that this pathway may be present in multiple *Microcystis* strains (bootstrap = 1, 97-99.6%). Similarity to enediyne biosynthesis proteins from marine cyanobacteria *Okeania* sp. was also observed (bootstrap = 0.855, ∼79% identity, Fig. 5).

**Figure 5:**
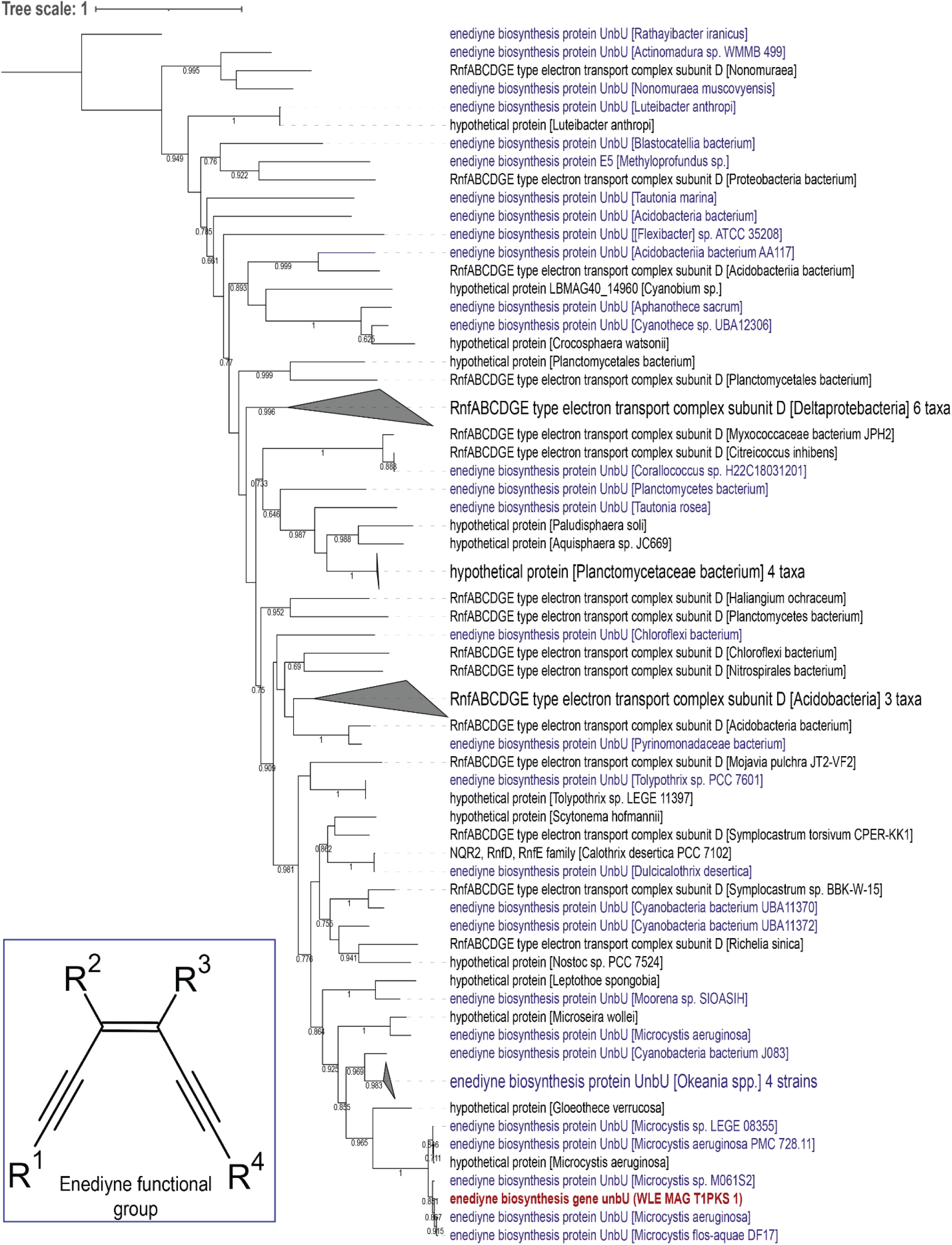
Protein phylogenies for A) a transport protein and B) core biosynthesis naringenin chalcone synthase from the T3PKS gene cluster. The transport protein has high homology to proteins involved in siderophore and cyclic peptide transport while the naringenin chalcone synthase has high homology to cyanobacterial naringenin chalcone synthases and some similarity to an alpha-pyrone synthesis polyketide synthase PKS-18 from the actinobacteria *Nocardia seriolae*.

Protein sequences from the T3PKS gene cluster also provided clues into potential functions of biosynthesis enzymes. An exporter protein sequence shared high sequence similarity to proteins involved in siderophore export from other *Microcysti*s strains (∼95-97% identity, bootstrap = 1, Fig S3A). Distantly related clades contained protein sequences with annotations for export of siderophores and/or cyclic peptides from a variety of cyanobacteria including benthic *Leptolyngbya* sp. (bootstrap = 0.894, ∼63% identity, Fig S3A). The core synthesis enzyme within this cluster, annotated by antiSMASH as naringenin-chalcone synthase, was most closely related to other *Microcystis* naringenin-chalcone synthase and T3PKS enzymes (∼98% identity, bootstrap = 0.877, Fig. S3B) as well as naringenin-chalcone synthases from *Chamaesiphon* sp. from the cyanobacteria Order *Synechococcales* (bootstrap = 0.688, 57-62% identity, Fig. S3B), none of which are linked to a known synthesized compound. Its protein sequence was also similar to an experimentally confirmed α-pyrone synthesis polyketide enzyme (PKS18) (Fig. 5B) from the actinobacterium *Nocarida seriolae* (50.6% identity, 97% alignment length).

### Transcriptional activity of BGCs

Quantification of transcript abundances revealed that BGC clusters were differentially expressed across stations and sampling times (Fig. 6). During early phases of the bloom, there was little expression of BGCs as seen on 21 July at WE2 and 29 July at WE4, likely due to low biomass of *Microcystis*. At WE12 during the peak bloom phase (4 August), clusters T3PKS, T1PKS 3, and micropeptin 3 had increased in relative transcript abundance compared to other clusters (Fig. 6). While the relative abundance of transcripts for *mcy* genes encoding microcystin (Yancey *et al*., 2022) was highest at WE12 during August, other BGCs had similar, and in some cases higher, transcript abundances than the microcystin encoding gene cluster during other phases of the bloom. The relative abundance of BGC transcripts shifted and greatly increased at all three stations during middle and late phases of the bloom, with high relative abundance of transcripts for T3PKS and T1PKS-NRPS-hgIEKS at WE12 on 25 August, WE2 on 6 October, and 8 September at WE4 (Fig. 6).

**Figure 6:**
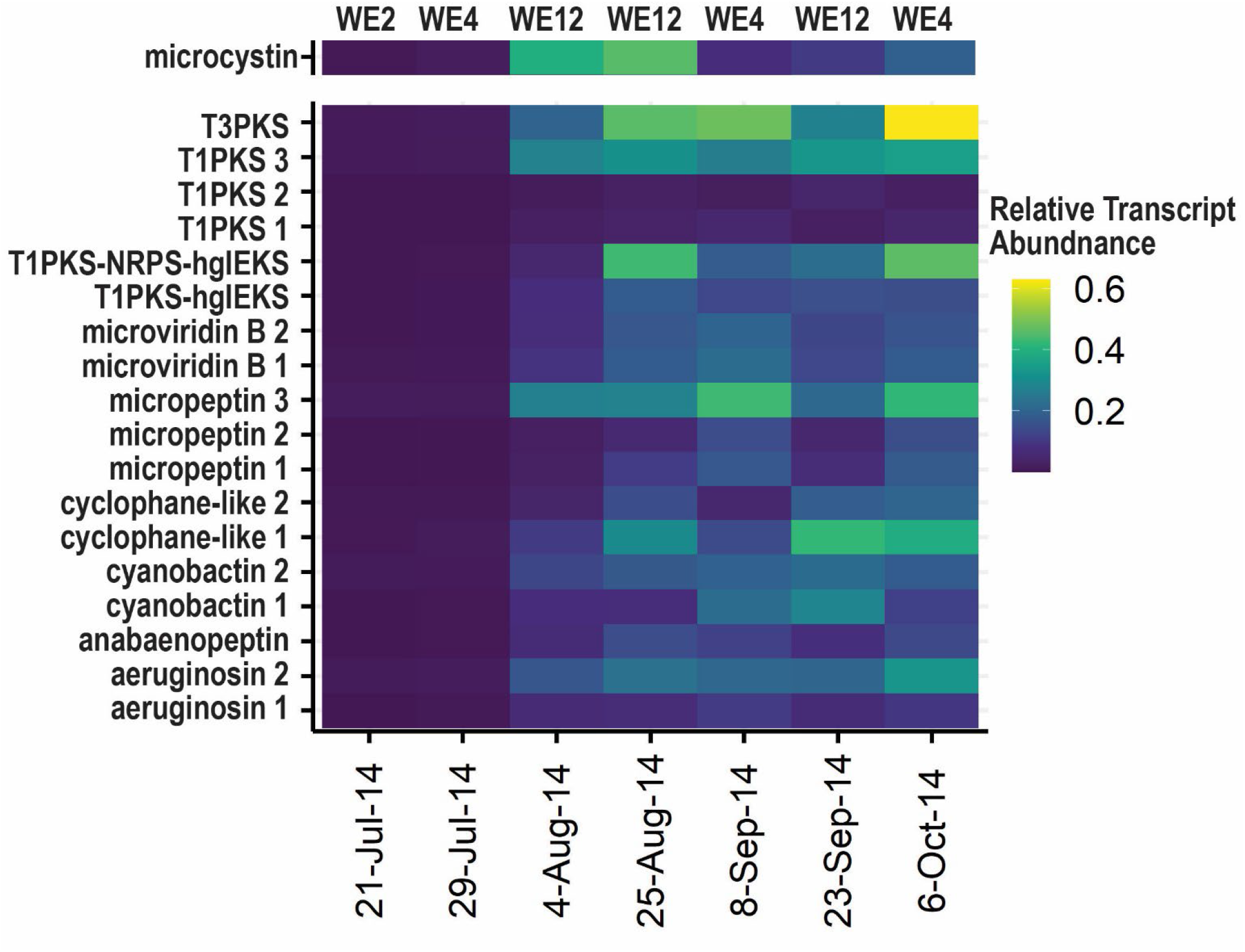
Relative abundance of transcripts from BGCs throughout the 2014 Bloom. Relative transcript abundance was calculated by determining the number of reads mapped to a BGC via specific cut off parameters, normalized to the number of reads mapped to an entire reference *Microcystis* genome to determine “expression effort” (see Experimental Procedure).

### Correlation Analysis between transcript abundance and both abiotic and biotic variables

To explore potential controls on the expression of BGCs, we performed correlational analysis of BGC transcript abundance with a variety of available abiotic conditions and relative abundances of known *Microcystis* predators and competitors in metagenomic data. Relative abundances of BGC transcripts were significantly correlated with both abiotic conditions (Table 2) and relative abundance of predatory and competitive organisms (Table 3). The transcript abundance of all BGC classes examined was significantly and negatively correlated with nitrate concentration while the transcript abundances of genes for cyanobactins (Pearson’s R=-0.531, p=0.029), cyclophane-like (Pearson’s R=0.598, p=0.040), and micropeptin (Pearson’s R=-0.412, p=0.089) BGCs were negatively correlated to temperature. Transcript abundances for aeruginosin, micropeptin, microviridin B, and cyclophane-like clusters had positive and significant or near significant correlations with both soluble reactive phosphorus (SRP) and total phosphorus (TP) concentrations (Table 2).

**Table 2:**
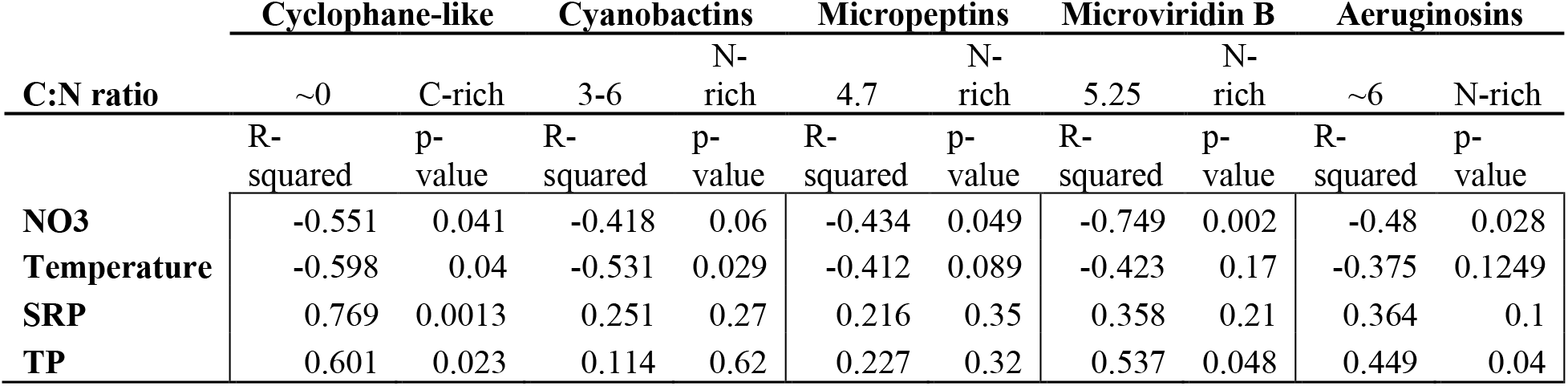
Abiotic and BGC expression Correlations.

**Table 3:** Pearson’s correlations for BGC expression and selected organism abundance

| Genus |  | aeruginosins |  | cyanobactins |  | micropeptins |  | microviridin B |  | cyclophane-like |  |
| --- | --- | --- | --- | --- | --- | --- | --- | --- | --- | --- | --- |
|  |  | R | p-value | R | p-value | R | p-value | R | p-value | R | p-value |
| Ciliates | *ICHTHYOPHTHIRIUS* | 0.553 | 0.033 | 0.245 | 0.378 | 0.336 | 0.221 | 0.653 | 0.041 | 0.799 | 0.006 |
|  | *PARAMECIUM* | 0.562 | 0.029 | 0.256 | 0.379 | 0.340 | 0.215 | 0.657 | 0.039 | 0.810 | 0.005 |
|  | *PSEUDOCOHNILEMBUS* | 0.549 | 0.034 | 0.230 | 0.410 | 0.326 | 0.236 | 0.641 | 0.046 | 0.803 | 0.005 |
|  | *TETRAHYMENA* | 0.563 | 0.029 | 0.238 | 0.393 | 0.335 | 0.222 | 0.659 | 0.038 | 0.817 | 0.004 |
| Diatoms | *CRASPEDOSTAUROS* | 0.567 | 0.054 | 0.298 | 0.347 | 0.436 | 0.156 | 0.414 | 0.308 | 0.853 | 0.007 |
|  | *SKELETONEMA* | 0.630 | 0.028 | 0.617 | 0.033 | 0.741 | 0.006 | 0.832 | 0.010 | 0.031 | 0.941 |
| Beta-proteobacteria | *PARABURKHOLDERIA* | 0.296 | 0.285 | 0.081 | 0.773 | 0.117 | 0.678 | 0.380 | 0.279 | 0.640 | 0.046 |
|  | *RHODOFERAX* | 0.409 | 0.066 | 0.419 | 0.059 | 0.488 | 0.025 | 0.669 | 0.009 | 0.171 | 0.559 |
| Proteobacteria | *OLIGOFLEXUS* | 0.505 | 0.094 | 0.057 | 0.860 | 0.191 | 0.553 | 0.530 | 0.177 | 0.828 | 0.011 |
| Cytophagia | *CHRYSEOTALEA* | 0.405 | 0.096 | 0.149 | 0.555 | 0.276 | 0.030 | 0.360 | 0.250 | 0.448 | 0.144 |

The relative metagenomic abundance of ciliates and diatoms exhibited some of the strongest, positive correlations with relative transcript abundance of BGCs, including those predicted for aeruginosin, microviridin B, and cyclophane-like clusters (Pearson’s R=0.653, p=0.041 to Pearson’s R=0.517, p=0.003, Table 3). Cyanobactin BGC expression was significantly and positively correlated with the relative abundance of the diatom *Skeletonema* (Pearson’s R= 0.617, p=0.033). Micropeptin BGC expression was also significantly and positively correlated with the abundance of *Skeletonema* (Pearson’s R= 0.741, p=0.006) as well as the betaproteobacterium, *Paraburkholderi*a (Pearson’s R= 0.488, p= 0.025). All other biotic correlations are summarized in Table S3.

## Discussion

*Microcystis* blooms are renowned for their production of microcystins, toxins that threaten drinking water supplies (Dawson, 1998; Campos and Vasconcelos, 2010; Bouaïcha et al., 2019). However, *Microcystis* genomes contain many other BGCs that encode known and unknown products, and their genetic diversity, biosynthesis, and ecological functions in the environment are poorly understood. In this study, we tracked spatial and temporal shifts in BGC abundance and transcriptional activity through seasonal changes in the biotic and abiotic environment of the 2014 western Lake Erie bloom, providing insights into the biosynthetic potential and expression of diverse *Microcystis* populations in their natural habitat. Our results demonstrate that *Microcystis* contains a diverse and highly dynamic suite of BGCs that encode both known cyanotoxins and cryptic compounds and are transcriptionally active in natural blooms. The transcriptional activity of these BGCs is related to both abiotic conditions (nutrients, temperature) as well as the composition of the biological community. While previous studies have identified biosynthetic gene clusters in *Microcystis* isolates (Humbert *et al*., 2013; Pérez-Carrascal *et al*., 2019; Pearson *et al*., 2020) or described metabolomic profiles in both field and culture studies (Janssen, 2019; le Manach *et al*., 2019), our work focused on the abundance, diversity, and drivers of transcriptional activity of BGCs within natural populations as they occur *in situ*.

*Microcystis* MAGs displayed extensive genetic diversity at the sub-species level, which is increasingly recognized to be prevalent in natural blooms (Berry *et al*., 2017; Smith *et al*., 2021; Yancey *et al*., 2022), and encodes important biosynthetic and physiological diversity (Steffen *et al*., 2017; Pérez-Carrascal *et al*., 2019; Yancey *et al*., 2022). Although these MAGs contained some redundancy and short contiguous sequences due to high strain heterogeneity present in natural populations (Dick *et al*., 2021; Yancey *et al*., 2022), they were sufficient to track spatial and temporal shifts in *Microcystis* gene content. Additionally, while our approach to estimating relative abundance of BGCs was based on normalization to *Microcystis* 16S rRNA abundance, and made the common assumption of one copy of the 16S rRNA gene per *Microcystis* genome (Vaitomaa *et al*., 2003; Davis *et al*., 2009), the nine closed genomes of *Microcystis* that are publicly available have two copies and there is uncertainty regarding the total copies of 16S rRNA genes in natural populations (Yancey *et al*., 2022). Regardless of these caveats, the results of this study do not depend on absolute calculations but rather relative differences, highlighting dynamics in time and space.

The diversity of BGCs we observed, and their spatiotemporal shift in abundance, reflects the presence and shifting abundance of diverse *Microcystis* strains with varying BGC content in their genomes. Whereas we identified and deeply analyzed nineteen BGCs, *Microcystis* typically has only 5-7 NRPS, PKS, or hybrid NRPS/PKS BGCs per genome (Shih *et al*., 2013; Pérez-Carrascal *et al*., 2019), and BGC content varies across clades (Pérez-Carrascal *et al*., 2019). Western Lake Erie *Microcystis* populations contain multiple strains in varying abundance (Rinta-Kanto *et al*., 2009; Smith *et al*., 2021; Yancey *et al*., 2022), often with microcystin producing strains dominating in early and peak bloom phases and non-microcystin producing strains dominating later phases of blooms (Berry *et al*., 2017; Yancey *et al*., 2022). Our results show that the “non-toxic” phases of the bloom, so-called because of low concentrations of microcystins, are enriched in other BGCs (Fig. 2) that are transcriptionally active, including anabaenopeptin, aeruginosin, micropeptin, and cyanobactins (Fig 6). Even more striking is the pervasive abundance and transcriptional activity of cryptic PKS-like BGCs with unknown products (Figs. 2,3,6). BGCs with the highest transcript abundance included those encoding unknown compounds such asT3PKS, T1PKS 3, cyclophane-like 1, and hybrid T1PKS-NRPS-hgIEKS (Fig. 6), indicating that they may be functionally and ecologically important. The lack of functional knowledge of these abundant and transcriptionally active PKSs highlights the need to investigate further their potential for generating unique metabolites.

The apparent “replacement” of *mcy* genes by other BGCs that we observed from early to late blooms in the field is consistent with studies of cultured *Microcystis* strains showing that non-microcystin producing strains often contain other BGCs in their place (Pérez-Carrascal *et al*., 2019). Our finding of a strong temporal shift in *Microcystis* BGC content and expression suggests environmental drivers of BGC content. Strains containing other BGCs instead of those encoding nitrogen-rich microcystins may be more fit during nitrogen deplete phases of the bloom that occur later in the season (Fig. 1B). For example, cyclophane-like molecules are nitrogen poor, in many cases lacking N atoms altogether (Table 2, (Martins *et al*., 2019)). Strains with BGCs that encode such molecules may have a competitive advantage over microcystin producers with higher N demands (van de Waal *et al*., 2009, 2014; Wagner *et al*., 2019). However, inconsistent with this conclusion is the increase in abundance of BGCs encoding other N-rich molecules, such as aeruginosin (Table 2), while nitrate concentrations decrease. These results suggest that the different secondary metabolites may have different fitness tradeoffs and/or environmental/physiological controls on gene expression. The high potential for biosynthesis of other secondary metabolites when microcystins are at low concentrations suggests functionally redundancy or niche differentiation among secondary metabolites.

BGCs that were more highly expressed in later bloom phases included anabaenopeptin, aeruginosin, micropeptin, and cyanobactin classes, which have inhibitory and toxic effects towards eukaryotes (Murakami *et al*., 1995; Welker and von Döhren, 2006b; Leikoski *et al*., 2012; Lenz *et al*., 2019). Other transcriptionally active BGCs encode unknown compounds with unknown toxicology. Given the continued emergence and discovery of novel cyanotoxins and their underpinning BGCs (Breinlinger *et al*., 2021; Lima *et al*., 2022), and the potential for toxicological synergism between multiple compounds (Fernandes *et al*., 2019; Pawlik-Skowrońska and Bownik, 2022), these results underscore the need to assess potential threats to human and ecosystem health. In addition to potential risks of unmonitored toxic cyanobacteria secondary metabolites, these molecules may be an untapped source for drug discovery (Aesoy and Herfindal, 2022). For example, the T3PKS BGC was one of the most abundant and transcriptionally active PKSs during early phases of the bloom and (Fig. 2,6) and contained a naringenin-chalcone synthase gene (Fig. 3), which may encode antibiotics or other compounds with wide ranges of biological activity and pharmaceutical relevance (Austin and Noel, 2003; Bhat *et al*., 2017). Predicted proteins from the T1PKS 3 BGC are similar to those found in biosynthesis of cytotoxic microginins, which have inhibitory properties against angiotensin-converting enzymes (Okino *et al*., 1993; Rounge *et al*., 2009). The T1PKS 1 (Fig 3A, Fig. S3A) cluster is predicted to encode biosynthesis of enediynes, which are excellent candidate antibiotics and anticancer compounds with rare structural characteristics (Nicolaou and Dai, 1991; Lanen and Shen, 2008). Finally, the presence of a tryptophan halogenase encoding gene with the NRPS-T1PKS-hgIEKS cluster (Fig. 3D) is noteworthy due to the broad pharmaceutical application of halogenated compounds, which have largely been studied in marine organisms up to this point (Gribble, 2015).

Correlation of transcript abundance with abiotic and biotic factors provided insights into environmental conditions that may influence expression of BGCs in natural environments. Negative correlations between relative abundance of transcripts and nitrate and temperature suggest a functional need for synthesized compounds in conditions of lower nitrate concentrations and temperature (Table 2). Positive correlations of BGC transcript abundance with abundances of known eukaryotic predators (Table 3) are consistent with the hypothesis that BGC products may deter grazers (ciliates) that commonly feed on cyanobacteria in aquatic systems (Calbet and Landry, 2004; Gobler *et al*., 2008). Of the cyanobacteria commonly found in western Lake Erie cyanoHABs, *Microcystis* are the most resistant to grazing by both daphnid and protozoan microorganisms (Ladds et al., 2021), consistent with findings that *Microcystis* is resistant to grazing due to production of secondary metabolites (Yang *et al*., 2006; Davis and Gobler, 2011; Sadler and von Elert, 2014). Likewise, significant correlations between expression of BGCs and abundance of photosynthetic competitors (diatoms) is consistent with a role of secondary metabolites in allelopathic interactions (Pflugmacher, 2002; Chia *et al*., 2018).

*Microcystis* secondary metabolites are likely multi-functional and may serve as nitrogen storage compounds (Harke and Gobler, 2013; Wang *et al*., 2021), aid in grazer defense (Wiegand and Pflugmacher, 2005; Penn *et al*., 2014; Sadler and von Elert, 2014), protection from reactive oxygen species (Alexova *et al*., 2011; Zilliges *et al*., 2011), and/or cell communication/phycosphere recruitment (Gan *et al*., 2012; Baran *et al*., 2015). While correlations uncovered here cannot determine true drivers of BGC abundance or expression, they provide hypotheses that can be tested with targeted experiments on the ecological and cellular role of the diversity of secondary metabolites encoded in *Microcystis* genomes.

## Conclusion

This study shows that natural populations of *Microcystis* within blooms contain and highly expresses a diverse and dynamic suite of genes encoding secondary metabolites. Several of these BGCs have no known products and are expressed most highly during the late phases of the bloom, which is dominated by “nontoxigenic” *Microcystis* strains that are incapable of producing microcystins. These results suggest the potential for diverse metabolite production beyond microcystins in western Lake Erie, with implications for understanding *Microcystis* physiology and diversity, community ecology, and water quality. Seasonal shifts in the abundance and expression of BGCs along with correlations with both abiotic and biotic variables suggest that the BGCs underpin adaptations to changing environmental conditions and may encode important niche differentiation among *Microcystis* strains. Several clusters identified encode toxic compounds while others encode cryptic compounds that may be toxic, highlighting the need to identify their products, determine their bioactivities, and assess potential threats to human and ecosystem health. Others show interesting chemical features that suggest avenues for exploring biotechnological and medicinal applications. Finally, the abundant environmental expression of BGCs encoding secondary metabolites that are of unknown function, and its correlation with biotic and abiotic conditions, suggest that the BGCs may play important roles in supporting the dominance of *Microcystis* over a wide range of conditions, underscoring the need to understand the functioning of these putative secondary metabolites in the ecophysiology and community interactions of *Microcystis* blooms.

## Acknowledgements

We thank Robert Hein, Teal Furnholm, and Derek Smith for the bioinformatic support. We also thank the field crew at CIGLR/GLERL including Paul Den Uyl, Dack Stuart, and Kent Baker for allowing us to sample with them and assisting in field measurements.

## Tables and Figure Captions

**Supplemental Figure 1:** Comparison of gene order, orientation, and sequence similarity between distinct BGCs from different *Microcystis* MAGs. Clinker was used to visually compare similarities and differences of the A) aeruginosin, B) cyanobactin, C) microviridin B, and D) micropeptin BGC classes. Homologous genes are the same color and percent similarity between genes is shown via a bar connecting homologous genes.

**Supplemental Figure 2:** Gene Schematics for identified BGC clusters including A) anabaenopeptin, B) microviridin B, C) aeruginosin, and D) a cyanobactin with 58% similarity to piricyclamide. Gene schematics were rendered through AntiSMASH v.6. Examples of chemical congeners that may be synthesized by these BGCs are shown.

**Supplemental Figure 3:** Phylogenies for A) a transport protein and B) core biosynthesis naringenin-chalcone synthase from the T3PKS gene cluster. The transport protein has high homology to proteins involved in siderophore transport while the naringenin chalcone synthase has is similar to cyanobacterial naringenin chalcone synthases. Bootstrap values are included at nodes where values are greater than 0.5. Proteins that share the same functional annotation of interest are colored. Dark red and bolded protein sequences are from the w. Lake Erie cyanoHAB as identified through antiSMASH.

**Supplemental Figure 4:** The gene schematic for microginin BGC identified in *Microcystis* aeruginosa LEGE 91341 deposited on the MiBIG database and described in Eusébio et al 2022. The schematic beneath it represents T1PKS 3 which shared 35% alignment and 93% identity to the complete microginin pathway shown above. The read mapping plot on the bottom demonstrates the presence of a microginin pathway in the 2014 w. Lake Erie cyanoHAB on 4-Augsut at WE2, with variable divergence from the reference sequence, suggesting this pathway is present in multiple cyanobacteria taxa.

## References

Aesoy, R. and Herfindal, L. (2022) Cyanobacterial anticancer compounds in clinical use: Lessons from the dolastatins and cryptophycins. The Pharmacological Potential of Cyanobacteria 55–79.

Alexova, R., Fujii, M., Birch, D., Cheng, J., Waite, T.D., Ferrari, B.C., and Neilan, B.A. (2011) Iron uptake and toxin synthesis in the bloom-forming Microcystis aeruginosa under iron limitation. Environ Microbiol 13: 1064–1077.

Allaire, J.J. (2015) RStudio: Integrated Development Environment for R, Boston.

Alneberg, J., Bjarnason, B.S., de Bruijn, I., Schirmer, M., Quick, J., Ijaz, U.Z., et al. (2013) CONCOCT: Clustering cONtigs on COverage and ComposiTion.

Altschul, S.F., Gish, W., Miller, W., Myers, E.W., and Lipman, D.J. (1990) Basic local alignment search tool. J Mol Biol 215: 403–410.

Arnold, J.B. (2021) ggthemes: Extra Themes, Scales and Geoms for “ggplot2.”

Austin, B., M. and Noel, J., P. (2003) The chalcone synthase superfamily of type III polyketide synthases. Nat Prod Rep 20: 79–110.

Bagchi, S.N., Sondhia, S., Agrawal, M.K., and Banerjee, S. (2016) An angiotensin-converting enzyme-inhibitory metabolite with partial structure of microginin in a cyanobacterium Anabaena fertilissima CCC597, producing fibrinolytic protease. J Appl Phycol 28: 177–180.

Baran, R., Brodie, E.L., Mayberry-Lewis, J., Hummel, E., da Rocha, U.N., Chakraborty, R., et al. (2015) Exometabolite niche partitioning among sympatric soil bacteria. Nature Communications 2015 6:1 6: 1–9.

Berry, M.A., Davis, T.W., Cory, R.M., Duhaime, M.B., Johengen, T.H., Kling, G.W., et al. (2017) Cyanobacterial harmful algal blooms are a biological disturbance to Western Lake Erie bacterial communities. Environ Microbiol 19: 1149–1162.

Beversdorf, L.J., Weirich, C.A., Bartlett, S.L., and Miller, T.R. (2017) Variable Cyanobacterial Toxin and Metabolite Profiles across Six Eutrophic Lakes of Differing Physiochemical Characteristics. Toxins 2017, Vol 9, Page 62 9: 62.

Bhat, Z.S., Rather, M.A., Maqbool, M., Lah, H.U., Yousuf, S.K., and Ahmad, Z. (2017) α-pyrones: Small molecules with versatile structural diversity reflected in multiple pharmacological activities-an update. Biomedicine & Pharmacotherapy 91: 265–277.

Blin, K., Shaw, S., Kloosterman, A.M., Charlop-Powers, Z., van Wezel, G.P., Medema, M.H., and Weber, T. (2021) antiSMASH 6.0: improving cluster detection and comparison capabilities. Nucleic Acids Res 49: W29–W35.

Bouaïcha, N., Miles, C.O., Beach, D.G., Labidi, Z., Djabri, A., Benayache, N.Y., and Nguyen-Quang, T. (2019) Structural diversity, characterization and toxicology of microcystins. Toxins (Basel) 11: 714.

Breinlinger, S., Phillips, T.J., Haram, B.N., Mareš, J., Martínez Yerena, J.A., Hrouzek, P., et al. (2021) Hunting the eagle killer: A cyanobacterial neurotoxin causes vacuolar myelinopathy. Science (1979) 371:.

Buchfink, B., Reuter, K., and Drost, H.G. (2021) Sensitive protein alignments at tree-of-life scale using DIAMOND. Nature Methods 2021 18:4 18: 366–368.

Bushnell, B. BBTools User Guide -DOE Joint Genome Institute.

Calbet, A. and Landry, M.R. (2004) Phytoplankton growth, microzooplankton grazing, and carbon cycling in marine systems. Limnol Oceanogr 49: 51–57.

Campos, A. and Vasconcelos, V. (2010) Molecular Mechanisms of Microcystin Toxicity in Animal Cells. International Journal of Molecular Sciences 2010, Vol 11, Pages 268-287 11: 268–287.

Carmichael, W.W. (1992) Cyanobacteria secondary metabolites-the cyanotoxins. Journal of Applied Bacteriology 72: 445–459.

Chapra, S.C., Boehlert, B., Fant, C., Victor J. Bierman, Jr., Henderson, J., Mills, D., et al. (2017) Climate Change Impacts on Harmful Algal Blooms in U.S. Freshwaters: A Screening-Level Assessment. Environ Sci Technol 51: 8933–8943.

Chen, L.X., Anantharaman, K., Shaiber, A., Murat Eren, A., and Banfield, J.F. (2020) Accurate and complete genomes from metagenomes. Genome Res 30: 315–333.

Chia, M.A., Jankowiak, J.G., Kramer, B.J., Goleski, J.A., Huang, I.S., Zimba, P. v., et al. (2018) Succession and toxicity of Microcystis and Anabaena (Dolichospermum) blooms are controlled by nutrient-dependent allelopathic interactions. Harmful Algae 74: 67–77.

Cooperative Institute for Great Lakes Research, U. of M.N.G.L.E.R.L. Physical, chemical, and biological water quality monitoring data to support detection of Harmful Algal Blooms (HABs) in western Lake Erie, collected by the Great Lakes Environmental Research Laboratory and the Cooperative Institute for Great Lakes Research since 2012.

Cragg, G.M. and Newman, D.J. (2013) Natural products: A continuing source of novel drug leads. Biochimica et Biophysica Acta (BBA) -General Subjects 1830: 3670–3695.

Davis, T.W., Berry, D.L., Boyer, G.L., and Gobler, C.J. (2009) The effects of temperature and nutrients on the growth and dynamics of toxic and non-toxic strains of Microcystis during cyanobacteria blooms. Harmful Algae 8: 715–725.

Davis, T.W. and Gobler, C.J. (2011) Grazing by mesozooplankton and microzooplankton on toxic and non-toxic strains of Microcystis in the Transquaking River, a tributary of Chesapeake Bay. J Plankton Res 33: 415–430.

Dawson, R.M. (1998) the toxicology of microcystins. Toxicon 36: 953–962.

Dick, G.J. (2018) Genomic Approaches in Earth and Environmental Sciences, 1st ed. John Wiley & Sons, Ltd.

Dick, G.J., Duhaime, M.B., Evans, J.T., Errera, R.M., Godwin, C.M., Kharbush, J.J., et al. (2021) The genetic and ecophysiological diversity of Microcystis. Environ Microbiol 1462-2920.15615.

Dittmann, E., Gugger, M., Sivonen, K., and Fewer, D.P. (2015) Natural Product Biosynthetic Diversity and Comparative Genomics of the Cyanobacteria. Trends Microbiol 23: 642–652.

Edgar, R.C. (2004) MUSCLE: multiple sequence alignment with high accuracy and high throughput. Nucleic Acids Res 32: 1792–1797.

Ehrenreich, I.M., Waterbury, J.B., and Webb, E.A. (2005) Distribution and diversity of natural product genes in marine and freshwater cyanobacterial cultures and genomes. Appl Environ Microbiol 71: 7401–13.

Eren, A.M., Esen, Ö.C., Quince, C., Vineis, J.H., Morrison, H.G., Sogin, M.L., and Delmont, T.O. (2015) Anvi’o: an advanced analysis and visualization platform for ‘omics data. PeerJ 3: e1319.

Eusébio, N., Castelo-Branco, R., Sousa, D., Preto, M., D’agostino, P., Gulder, T.A.M., and Leão, P.N. (2022) Discovery and heterologous expression of microginins from Microcystis aeruginosa LEGE 91341.

Fernandes, K., Gomes, A., Calado, L., Yasui, G., Assis, D., Henry, T., et al. (2019) Toxicity of Cyanopeptides from Two Microcystis Strains on Larval Development of Astyanax altiparanae. Toxins (Basel) 11:.

Gan, N., Xiao, Y., Zhu, L., Wu, Z., Liu, J., Hu, C., and Song, L. (2012) The role of microcystins in maintaining colonies of bloom-forming Microcystis spp. Environ Microbiol 14: 730–742.

Gilchrist, C.L.M. and Chooi, Y.-H. (2021) clinker & clustermap.js: automatic generation of gene cluster comparison figures. Bioinformatics.

Gobler, C.J., Davis, T.W., Deonarine, S.N., Saxton, M.A., Lavrentyev, P.J., Jochem, F.J., and Wilhelm, S.W. (2008) Grazing and virus-induced mortality of microbial populations before and during the onset of annual hypoxia in Lake Erie. Aquatic Microbial Ecology 51: 117–128.

Gribble, G.W. (2015) Biological Activity of Recently Discovered Halogenated Marine Natural Products. Marine Drugs 2015, Vol 13, Pages 4044–4136 13: 4044–4136.

Griffith, A.W. and Gobler, C.J. (2020) Harmful algal blooms: A climate change co-stressor in marine and freshwater ecosystems. Harmful Algae 91: 101590.

Harke, M.J. and Gobler, C.J. (2013) Global Transcriptional Responses of the Toxic Cyanobacterium, Microcystis aeruginosa, to Nitrogen Stress, Phosphorus Stress, and Growth on Organic Matter. PLoS One 8: e69834.

Harke, M.J., Steffen, M.M., Otten, T.G., Wilhelm, S.W., Wood, S.A., and Paerl, H.W. (2016) A review of the global ecology, genomics, and biogeography of the toxic cyanobacterium, Microcystis spp. Harmful Algae 54: 4–20.

Heisler, J., Glibert, P.M., Burkholder, J.M., Anderson, D.M., Cochlan, W., Dennison, W.C., et al. (2008) Eutrophication and harmful algal blooms: A scientific consensus. Harmful Algae 8: 3–13.

Ho, J.C., Michalak, A.M., and Pahlevan, N. (2019) Widespread global increase in intense lake phytoplankton blooms since the 1980s. Nature 574: 667–670.

Hughes, E.O., Gorham, P.R., and Zehnder, A. (1958) Toxicity of a unialgal culture of Microcystis aeruginosa. Can J Microbiol 4: 225–236.

Huisman, J., Codd, G.A., Paerl, H.W., Ibelings, B.W., H Verspagen, J.M., and Visser, P.M. (2018) Cyanobacterial blooms. Nat Rev Microbiol.

Humbert, J.-F., Barbe, V., Latifi, A., Gugger, M., Calteau, A., Coursin, T., et al. (2013) A Tribute to Disorder in the Genome of the Bloom-Forming Freshwater Cyanobacterium Microcystis aeruginosa. PLoS One 8: e70747.

Janssen, E.M.L. (2019) Cyanobacterial peptides beyond microcystins – A review on co-occurrence, toxicity, and challenges for risk assessment. Water Res 151: 488–499.

Kang, D.D., Froula, J., Egan, R., and Wang, Z. (2015) MetaBAT, an efficient tool for accurately reconstructing single genomes from complex microbial communities. PeerJ 3: e1165.

Kassambara, A. (2020) ggpubr: “ggplot2” Based Publication Ready Plots.

Kehr, J.-C., Gatte Picchi, D., and Dittmann, E. (2011) Natural product biosyntheses in cyanobacteria: A treasure trove of unique enzymes. Beilstein journal of organic chemistry 7: 1622–35.

Kotut, K., Ballot, A., and Krienitz, L. (2006) Toxic cyanobacteria and their toxins in standing waters of Kenya: implications for water resource use. J Water Health 4: 233–245.

Krausfeldt, L.E., Farmer, A.T., Castro Gonzalez, H.F., Zepernick, B.N., Campagna, S.R., and Wilhelm, S.W. (2019) Urea Is Both a Carbon and Nitrogen Source for Microcystis aeruginosa: Tracking 13C Incorporation at Bloom pH Conditions. Front Microbiol 10: 1064.

Kust, A., Řeháková, K., Vrba, J., Maicher, V., Mareš, J., Hrouzek, P., et al. (2020) Insight into Unprecedented Diversity of Cyanopeptides in Eutrophic Ponds Using an MS/MS Networking Approach. Toxins 2020, Vol 12, Page 561 12: 561.

Laczny, C.C., Sternal, T., Plugaru, V., Gawron, P., Atashpendar, A., Margossian, H., et al. (2015) VizBin -an application for reference-independent visualization and human-augmented binning of metagenomic data. Microbiome 3: 1.

Ladds, M., Jankowiak, J., and Gobler, C.J. (2021) Novel high throughput sequencing -fluorometric approach demonstrates Microcystis blooms across western Lake Erie are promoted by grazing resistance and nutrient enhanced growth. Harmful Algae 110: 102126.

Lake Erie Programs | Ohio Environmental Protection Agency.

Lanen, S.G. van and Shen, B. (2008) Biosynthesis of Enediyne Antitumor Antibiotics. Curr Top Med Chem 8: 448.

Langmead, B. and Salzberg, S.L. (2012) Fast gapped-read alignment with Bowtie 2. Nat Methods 9: 357–359.

Leikoski, N., Fewer, D.P., Jokela, J., Alakoski, P., Wahlsten, M., and Sivonen, K. (2012) Analysis of an Inactive Cyanobactin Biosynthetic Gene Cluster Leads to Discovery of New Natural Products from Strains of the Genus Microcystis. PLoS One 7: e43002.

Lenz, K.A., Miller, T.R., and Ma, H. (2019) Anabaenopeptins and cyanopeptolins induce systemic toxicity effects in a model organism the nematode Caenorhabditis elegans. Chemosphere 214: 60–69.

Li, D., Liu, C.-M., Luo, R., Sadakane, K., and Lam, T.-W. (2015) MEGAHIT: an ultra-fast single-node solution for large and complex metagenomics assembly via succinct de Bruijn graph. Bioinformatics 31: 1674–1676.

Lima, S.T., Fallon, T.R., Cordoza, J.L., Chekan, J.R., Delbaje, E., Hopiavuori, A.R., et al. (2022) Biosynthesis of Guanitoxin Enables Global Environmental Detection in Freshwater Cyanobacteria. J Am Chem Soc 144: 9372–9379.

Madden, T. (2013) The BLAST Sequence Analysis Tool.

le Manach, S., Duval, C., Marie, A., Djediat, C., Catherine, A., Edery, M., et al. (2019) Global Metabolomic Characterizations of Microcystis spp. Highlights Clonal Diversity in Natural Bloom-Forming Populations and Expands Metabolite Structural Diversity. Front Microbiol 10: 791.

Martins, T.P., Rouger, C., Glasser, N.R., Freitas, S., de Fraissinette, N.B., Balskus, E.P., et al. (2019) Chemistry, bioactivity and biosynthesis of cyanobacterial alkylresorcinols. Nat Prod Rep 36: 1437–1461.

Murakami, M., Ishida, K., Okino, T., Okita, Y., Matsuda, H., and Yamaguchi, K. (1995) Aeruginosins 98-A and B, trypsin inhibitors from the blue-green alga Microcystis aeruginosa (NIES-98). Tetrahedron Lett 36: 2785–2788.

Nicolaou, K.C. and Dai, W.-M (1991) Chemistry and Biology of the Enediyne Anticancer Antibiotics. Angewandte Chemie International Edition in English 30: 1387–1416.

Okino, T., Matsuda, H., Murakami, M., and Yamaguchi, K. (1993) Microginin, an angiotensin-converting enzyme inhibitor from the blue-green alga Microcystis aeruginosa. Tetrahedron Lett 34: 501–504.

Paerl, H.W., Otten, T.G., and Kudela, R. (2018) Mitigating the Expansion of Harmful Algal Blooms Across the Freshwater-to-Marine Continuum. Environ Sci Technol 52: 5519–5529.

Parks, D.H., Imelfort, M., Skennerton, C.T., Hugenholtz, P., and Tyson, G.W. (2015) CheckM: assessing the quality of microbial genomes recovered from isolates, single cells, and metagenomes. Genome Res 25: 1043–55.

Pawlik-Skowrońska, B. and Bownik, A. (2022) Synergistic toxicity of some cyanobacterial oligopeptides to physiological activities of Daphnia magna (Crustacea). Toxicon 206: 74–84.

Pearson, L.A., Crosbie, N.D., and Neilan, B.A. (2020) Distribution and conservation of known secondary metabolite biosynthesis gene clusters in the genomes of geographically diverse Microcystis aeruginosa strains. Mar Freshw Res 71: 701.

Pearson, L.A., Dittmann, E., Mazmouz, R., Ongley, S.E., D’Agostino, P.M., and Neilan, B.A. (2016) The genetics, biosynthesis and regulation of toxic specialized metabolites of cyanobacteria. Harmful Algae 54: 98–111.

Penn, K., Wang, J., Fernando, S.C., and Thompson, J.R. (2014) Secondary metabolite gene expression and interplay of bacterial functions in a tropical freshwater cyanobacterial bloom. ISME J 8: 1866–1878.

Pérez-Carrascal, O.M., Terrat, Y., Giani, A., Fortin, N., Greer, C.W., Tromas, N., and Shapiro, B.J. (2019) Coherence of Microcystis species revealed through population genomics. bioRxiv 541755.

Pflugmacher, S. (2002) Possible allelopathic effects of cyanotoxins, with reference to microcystin-LR, in aquatic ecosystems. Environ Toxicol 17: 407–413.

Price, M.N., Dehal, P.S., and Arkin, A.P. (2009) FastTree: Computing Large Minimum Evolution Trees with Profiles instead of a Distance Matrix. Mol Biol Evol 26: 1641–1650.

QGIS Development Team (2020) QGIS Geo Graphic Information System. Open Source Geospatial 742 Foundation Project.

Qin, B., Zhu, G., Gao, G., Zhang, Y., Li, W., Paerl, H.W., and Carmichael, W.W. (2010) A Drinking Water Crisis in Lake Taihu, China: Linkage to Climatic Variability and Lake Management. Environ Manage 45: 105–112.

Quast, C., Pruesse, E., Yilmaz, P., Gerken, J., Schweer, T., Yarza, P., et al. (2013) The SILVA ribosomal RNA gene database project: Improved data processing and web-based tools. Nucleic Acids Res 41: D590–D596.

Rinta-Kanto, J.M., Konopko, E.A., DeBruyn, J.M., Bourbonniere, R.A., Boyer, G.L., and Wilhelm, S.W. (2009) Lake Erie Microcystis: Relationship between microcystin production, dynamics of genotypes and environmental parameters in a large lake. Harmful Algae 8: 665–673.

Rounge, T.B., Rohrlack, T., Nederbragt, A.J., Kristensen, T., and Jakobsen, K.S. (2009) A genome-wide analysis of nonribosomal peptide synthetase gene clusters and their peptides in a Planktothrix rubescens strain. BMC Genomics 10: 396.

Sadler, T. and von Elert, E. (2014) Physiological interaction of Daphnia and Microcystis with regard to cyanobacterial secondary metabolites. Aquatic Toxicology 156: 96–105.

Shih, P.M., Wu, D., Latifi, A., Axen, S.D., Fewer, D.P., Talla, E., et al. (2013) Improving the coverage of the cyanobacterial phylum using diversity-driven genome sequencing. Proc Natl Acad Sci U S A 110: 1053–1058.

Sieber, C.M.K., Probst, A.J., Sharrar, A., Thomas, B.C., Hess, M., Tringe, S.G., and Banfield, J.F. (2018) Recovery of genomes from metagenomes via a dereplication, aggregation and scoring strategy. Nat Microbiol 3: 836–843.

Skinnider, M.A., Johnston, C.W., Gunabalasingam, M., Merwin, N.J., Kieliszek, A.M., MacLellan, R.J., et al. (2020) Comprehensive prediction of secondary metabolite structure and biological activity from microbial genome sequences. Nat Commun 11: 1–9.

Smith, D.J., Tan, J.Y., Powers, M.A., Lin, X.N., Davis, T.W., and Dick, G.J. (2021) Individual Microcystis colonies harbour distinct bacterial communities that differ by Microcystis oligotype and with time. Environ Microbiol 23: 3020–3036.

Steffen, M.M., Belisle, B.S., Watson, S.B., Boyer, G.L., and Wilhelm, S.W. (2014) Status, causes and controls of cyanobacterial blooms in Lake Erie. J Great Lakes Res 40: 215–225.

Steffen, M.M., Davis, T.W., McKay, R.M.L., Bullerjahn, G.S., Krausfeldt, L.E., Stough, J.M.A., et al. (2017) Ecophysiological Examination of the Lake Erie Microcystis Bloom in 2014: Linkages between Biology and the Water Supply Shutdown of Toledo, OH. Environ Sci Technol 51: 6745–6755.

Stewart, I., Seawright, A.A., and Shaw, G.R. (2008) Cyanobacterial poisoning in livestock, wild mammals and birds--an overview. Adv Exp Med Biol 619: 613–637.

Tillett, D., Dittmann, E., Erhard, M., Von Döhren, H., Börner, T., and Neilan, B.A. (2000) Structural organization of microcystin biosynthesis in Microcystis aeruginosa PCC7806: An integrated peptide-polyketide synthetase system. Chem Biol 7: 753–764.

Ultsch, A. and Mörchen, F. (2009) ESOM-Maps: tools for clustering, visualization, and classification with Emergent SOM Analysis of high dimensional and big data View project Genetic Foundations of Leukemia View project ESOM-Maps: tools for clustering, visualization, and classification with Emergent SOM.

Vaitomaa, J., Rantala, A., Halinen, K., Rouhiainen, L., Tallberg, P., Mokelke, L., and Sivonen, K. (2003) Quantitative real-time PCR for determination of microcystin synthetase e copy numbers for microcystis and anabaena in lakes. Appl Environ Microbiol 69: 7289–7297.

van de Waal, D., Smith, V., Declerck, S., Stam, E., and Esler, J. (2014) Stoichiometric regulation of phytoplankton toxins. Ecol Lett 17: 736–742.

van de Waal, D.B., Verspagen, J.M.H., Lürling, M., van Donk, E., Visser, P.M., and Huisman, J. (2009) The ecological stoichiometry of toxins produced by harmful cyanobacteria: An experimental test of the carbon-nutrient balance hypothesis. Ecol Lett 12: 1326–1335.

Wagner, N.D., Osburn, F.S., Wang, J., Taylor, R.B., Boedecker, A.R., Chambliss, C.K., et al. (2019) Biological stoichiometry regulates toxin production in microcystis aeruginosa (UTEX 2385). Toxins (Basel) 11: 601.

Wang, J., Wagner, N.D., Fulton, J.M., and Scott, J.T. (2021) Diazotrophs modulate phycobiliproteins and nitrogen stoichiometry differently than other cyanobacteria in response to light and nitrogen availability. Limnol Oceanogr 66: 2333–2345.

Watson, S.B., Miller, C., Arhonditsis, G., Boyer, G.L., Carmichael, W., Charlton, M.N., et al. (2016) The re-eutrophication of Lake Erie: Harmful algal blooms and hypoxia. Harmful Algae 56: 44–66.

Welker, M. and von Döhren, H. (2006) Cyanobacterial peptides -Nature’s own combinatorial biosynthesis. FEMS Microbiol Rev 30: 530–563.

Wick, R.R., Schultz, M.B., Zobel, J., and Holt, K.E. (2015) Bandage: interactive visualization of de novo genome assemblies: Fig. 1. Bioinformatics 31: 3350–3352.

Wickham, H. (2011) ggplot2. Wiley Interdiscip Rev Comput Stat 3: 180–185.

Wiegand, C. and Pflugmacher, S. (2005) Ecotoxicological effects of selected cyanobacterial secondary metabolites a short review. Toxicol Appl Pharmacol 203: 201–218.

Yancey, C.E., Smith, D.J., Uyl, P.A. den, Mohamed, O.G., Yu, F., Ruberg, S.A., et al. (2022) Metagenomic and Metatranscriptomic Insights into Population Diversity of Microcystis Blooms: Spatial and Temporal Dynamics of mcy Genotypes, Including a Partial Operon That Can Be Abundant and Expressed. Appl Environ Microbiol.

Yang, Z., Kong, F., Shi, X., and Cao, H. (2006) Morphological Response of Microcystis aeruginosa to Grazing by Different Sorts of Zooplankton. Hydrobiologia 2006 563:1 563: 225–230.

Zilliges, Y., Kehr, J.C., Meissner, S., Ishida, K., Mikkat, S., Hagemann, M., et al. (2011) The Cyanobacterial Hepatotoxin Microcystin Binds to Proteins and Increases the Fitness of Microcystis under Oxidative Stress Conditions. PLoS One 6: e17615.

