## Supplementary material for "*Expression of* Microcystis *biosynthetic gene clusters in natural populations suggests temporally dynamic synthesis of novel and known secondary metabolites in western Lake Erie*": Fig. S

**SUPPLEMENTALS**

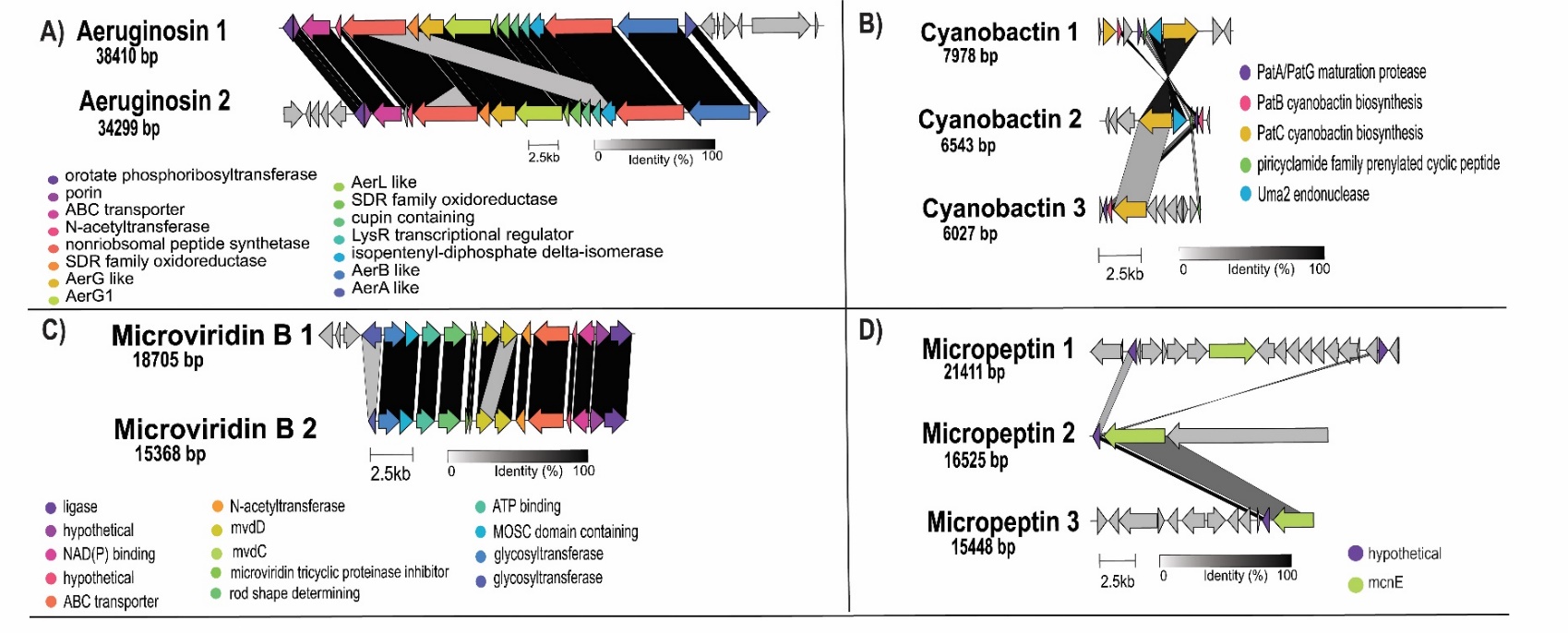
**Supplemental Figure 1:** Comparison of gene order, orientation, and sequence similarity between distinct BGCs from different *Microcystis* MAGs. Clinker was used to visually compare similarities and differences of the A) aeruginosin, B) cyanobactin, C) microviridin B, and D) micropeptin BGC classes. Homologous genes are the same color and percent similarity between genes is shown via a bar connecting homologous genes.

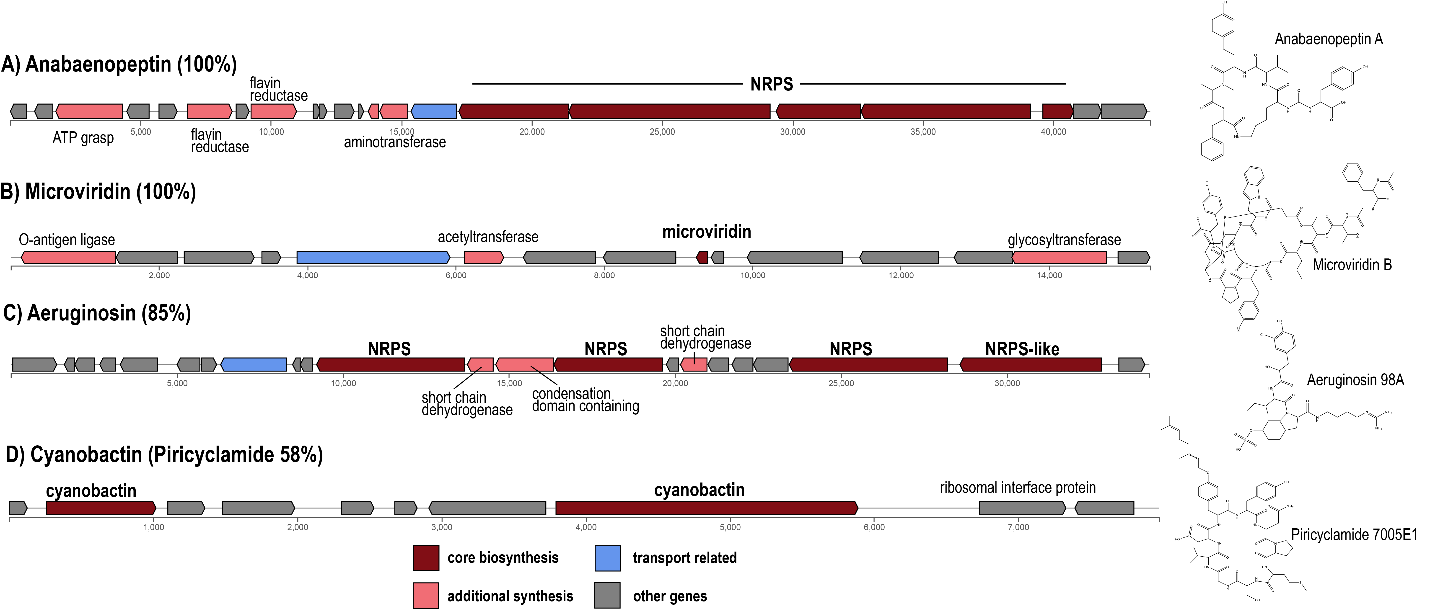

**Supplemental Figure 2:** Gene Schematics for identified BGC clusters including A) anabaenopeptin, B) microviridin B, C) aeruginosin, and D) a cyanobactin with 58% similarity to piricyclamide. Gene schematics were rendered through AntiSMASH v.6. Examples of chemical congeners that may be synthesized by these BGCs are shown. Percent similarity represents the percentage of genes within the closest known compound on the MiBIG database that have a significant BLAST hit

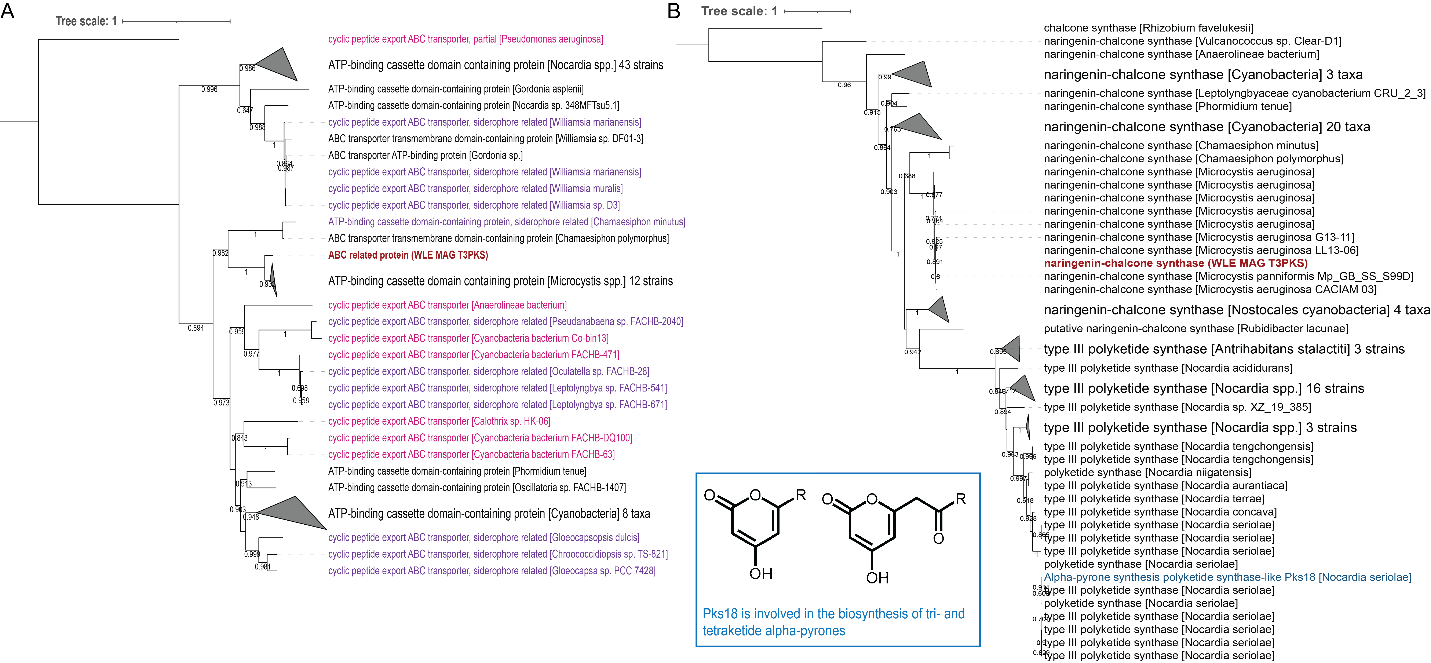

**Supplemental Figure 3:** Phylogenies for A) a transport protein and B) core biosynthesis naringenin-chalcone synthase from the T3PKS gene cluster. The transport protein has high homology to proteins involved in siderophore transport while the naringenin chalcone synthase has is similar to cyanobacterial naringenin chalcone synthases*.* Bootstrap values are included at nodes where values are greater than 0.5. Proteins that share the same functional annotation of interest are colored. Dark red and bolded protein sequences are from the W. Lake Erie cyanoHAB as identified through antiSMASH.

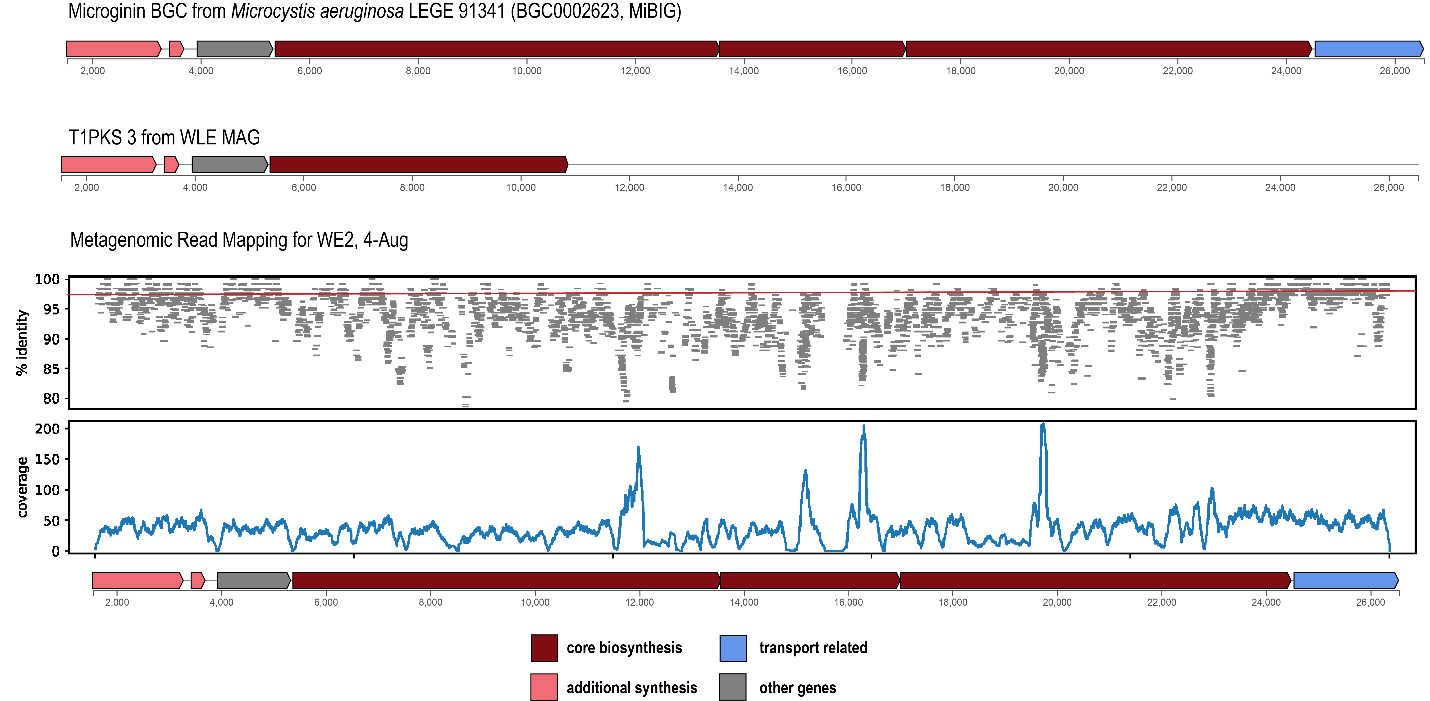

**Supplemental Figure 4:** The gene schematic for microginin BGC identified in *Microcystis aeruginosa* LEGE 91341 deposited on the MiBIG database and described in Eusébio et al 2022. The schematic beneath it represents T1PKS 3 which shared 35% alignment and 93% identity to the complete microginin pathway shown above. The read mapping plot on the bottom demonstrates the presence of a microginin pathway in the 2014 w. Lake Erie cyanoHAB on 4-Augsut at WE2, with variable divergence from the reference sequence, suggesting this pathway is present in multiple cyanobacteria taxa.

**Table S1:** PKS Cluster Similarities to known compounds as identified via antiSMASH.

| Cluster ID | Known Compound Cluster Hit | Similarity (%) |
| --- | --- | --- |
| T1PKS 1 | Calicheamicin | 4 |
| T1PKS 2 | Carbamidocyclophane | 30 |
| T1PKS 3 | Cylindrocyclophane | 40 |
| T3PKS | NA | NA |
| T1PKS-NRPS-hgIEKS | Phenalamide | 40 |
| T1PKS-hgIEKS | Bartoloside | 27 |

**Table S2**: Correlations of BGC relative transcript abundance and available abiotic variables

| **Cyanobactin** |  | |
| --- | --- | --- |
| Variable | R-squared | p-value |
| NO3 | -0.418 | 0.59 |
| NH4 | 0.188 | 0.41 |
| Temperature | -0.514 | 0.029 |
| pH | -0.129 | 0.58 |
| SRP | 0.251 | 0.27 |
| TP | 0.114 | 0.62 |
| PC | -0.0536 | 0.82 |
| chlA | -0.0603 | 0.8 |
| **Aeruginosins** |  | |
| Variable | R-squared | p-value |
| NO3 | -0.48 | 0.028 |
| NH4 | 0.0988 | 0.67 |
| Temperature | -0.375 | 0.1249 |
| pH | 0.0413 | 0.859 |
| SRP | 0.364 | 0.1 |
| TP | 0.449 | 0.04 |
| PC | 0.252 | 0.27 |
| chlA | 0.162 | 0.48 |
| **Cyclophane-like** | | |
| Variable | R-squared | p-value |
| NO3 | -0.551 | 0.041 |
| NH4 | 0.0802 | 0.79 |
| Temperature | -0.598 | 0.04 |
| pH | -0.238 | 0.41 |
| SRP | 0.769 | 0.0013 |
| TP | 0.601 | 0.023 |
| PC | 0.129 | 0.66 |
| chlA | -0.0753 | 0.8 |
| **Micropeptins** |  | |
| Variable | R-squared | p-value |
| NO3 | -0.434 | 0.049 |
| NH4 | 0.178 | 0.44 |
| Temperature | -0.412 | 0.089 |
| pH | -0.00607 | 0.98 |
| SRP | 0.216 | 0.35 |
| TP | 0.227 | 0.32 |
| PC | 0.0991 | 0.669 |
| chlA | 0.0839 | 0.72 |
| **Microviridin B** | | |
| Variable | R-squared | p-value |
| NO3 | -0.749 | 0.002 |
| NH4 | 0.00647 | 0.98 |
| Temperature | -0.423 | 0.17 |
| pH | 0.144 | 0.62 |
| SRP | 0.358 | 0.21 |
| TP | 0.537 | 0.048 |
| PC | 0.312 | 0.28 |
| chlA | 0.218 | 0.45 |

**Table S3:** Correlations of BGC relative transcript abundance and relative abundance of select organisms

| **BGC Class** | **Organism Genus** | | | **R-squared** | | **p-value** |
| --- | --- | --- | --- | --- | --- | --- |
| Cyanobactin | *UNCLASSIFIED_GEMMATIMONADACEAE_GENUS* | | | 0.532002 | | 0.023055 |
| Cyanobactin | *HERBASPIRILLUM* | | | -0.64425 | | 0.023743 |
| Cyanobactin | *WOLBACHIA* | | | -0.52319 | | 0.025879 |
| Cyanobactin | *UNCLASSIFIED_CYTOPHAGALES_GENUS* | | | 0.444163 | | 0.04368 |
| Cyanobactin | *RHODOFERAX* | | | 0.418508 | | 0.059008 |
| Cyanobactin | *GEMMATIMONAS* | | | -0.41654 | | 0.060332 |
| Cyanobactin | *ROSEOMONAS* | | | -0.4043 | | 0.069096 |
| Cyanobactin | *UNCLASSIFIED_VERRUCOMICROBIACEAE_GENUS* | | | -0.45766 | | 0.086274 |
| Cyanobactin | *UNCLASSIFIED_ARMATIMONADETES_GENUS* | | | -0.378 | | 0.09112 |
| Cyanobactin | *ACINETOBACTER* | | | -0.50547 | | 0.093656 |
| Cyanobactin | *NOVIHERBASPIRILLUM* | | | -0.48984 | | 0.105981 |
| Cyanobactin | *UNCLASSIFIED_GEMMATIMONADETES_GENUS* | | | 0.352644 | | 0.116898 |
| Cyanobactin | *UNCLASSIFIED_SYNECHOCOCCALES_GENUS* | | | -0.41704 | | 0.121975 |
| Cyanobactin | *BREVUNDIMONAS* | | | -0.34195 | | 0.129211 |
| Cyanobactin | *PIRELLULA* | | | -0.35852 | | 0.144021 |
| Cyanobactin | *BDELLOVIBRIO* | | | -0.35723 | | 0.145583 |
| Cyanobactin | *UNCLASSIFIED_PHYCISPHAERAE_GENUS* | | | -0.35563 | | 0.147523 |
| Cyanobactin | *SKELETONEMA* | | | 0.617171 | | 0.03252 |
| Cyanobactin | *APOSTICHOPUS* | | | -0.51817 | | 0.08438 |
| Cyanobactin | *PHYTOPHTHORA* | | | 0.372823 | | 0.12757 |
| Cyanobactin | *EURYTEMORA* | | | 0.32275 | | 0.153585 |
| Aer/Ana | *UNCLASSIFIED_BATHYARCHAEOTA_GENUS* | | | -0.54321 | | 0.010932 |
| Aer/Ana | *UNCLASSIFIED_EURYARCHAEOTA_GENUS* | | | -0.5217 | | 0.081919 |
| Aer/Ana | *WOLBACHIA* | | | -0.68302 | | 0.001783 |
| Aer/Ana | *UNCLASSIFIED_VERRUCOMICROBIACEAE_GENUS* | | | -0.71213 | | 0.002897 |
| Aer/Ana | *GEMMATIMONAS* | | | -0.56891 | | 0.007115 |
| Aer/Ana | *ACINETOBACTER* | | | -0.72704 | | 0.007383 |
| Aer/Ana | *PAUCIBACTER* | | | -0.71693 | | 0.008691 |
| Aer/Ana | *UNCLASSIFIED_ARMATIMONADETES_GENUS* | | | -0.54466 | | 0.010681 |
| Aer/Ana | *UNCLASSIFIED_SYNECHOCOCCALES_GENUS* | | | -0.62251 | | 0.013194 |
| Aer/Ana | *UNCLASSIFIED_CHLOROFLEXI_GENUS* | | | -0.50549 | | 0.019408 |
| Aer/Ana | *BRYOBACTER* | | | -0.58735 | | 0.021321 |
| Aer/Ana | *UNCLASSIFIED_VERRUCOMICROBIALES_GENUS* | | | -0.49482 | | 0.02258 |
| Aer/Ana | *UNCLASSIFIED_VERRUCOMICROBIA_GENUS* | | | -0.49006 | | 0.024122 |
| Aer/Ana | *ROSEOMONAS* | | | -0.48363 | | 0.026336 |
| Aer/Ana | *POLYNUCLEOBACTER* | | | -0.56913 | | 0.02681 |
| Aer/Ana | *METHYLOBACTERIUM* | | | -0.47444 | | 0.029776 |
| Aer/Ana | *UNCLASSIFIED_ACTINOMYCETIA_GENUS* | | | -0.47312 | | 0.0303 |
| Aer/Ana | *RUNELLA* | | | -0.62123 | | 0.031077 |
| Aer/Ana | *UNCLASSIFIED_GEMMATIMONADACEAE_GENUS* | | | 0.497769 | | 0.035545 |
| Aer/Ana | *UNCLASSIFIED_RHODOBACTERACEAE_GENUS* | | | -0.49136 | | 0.03837 |
| Aer/Ana | *SYNECHOCOCCUS* | | | -0.53619 | | 0.03936 |
| Aer/Ana | *LIMNOHABITANS* | | | -0.43737 | | 0.047397 |
| Aer/Ana | *UNCLASSIFIED_OPITUTAE_GENUS* | | | -0.51859 | | 0.047632 |
| Aer/Ana | *GEMMOBACTER* | | | -0.43359 | | 0.049571 |
| Aer/Ana | *BREVUNDIMONAS* | | | -0.43254 | | 0.050189 |
| Aer/Ana | *UNCLASSIFIED_FLAVOBACTERIACEAE_GENUS* | | | -0.50918 | | 0.052554 |
| Aer/Ana | *HERBASPIRILLUM* | | | -0.5664 | | 0.054859 |
| Aer/Ana | *PLANKTOPHILA* | | | -0.55568 | | 0.060676 |
| Aer/Ana | *UNCLASSIFIED_PELAGIBACTERALES_GENUS* | | | -0.54773 | | 0.065257 |
| Aer/Ana | *RHODOFERAX* | | | 0.408822 | | 0.065753 |
| Aer/Ana | *BELNAPIA* | | | -0.48663 | | 0.065843 |
| Aer/Ana | *UNCLASSIFIED_BRYOBACTERALES_GENUS* | | | -0.46307 | | 0.082158 |
| Aer/Ana | *TABRIZICOLA* | | | -0.37622 | | 0.092781 |
| Aer/Ana | *OLIGOFLEXUS* | | | 0.504783 | | 0.094177 |
| Aer/Ana | *CHRYSEOTALEA* | | | 0.404708 | | 0.095728 |
| Aer/Ana | *UNCLASSIFIED_PHYCISPHAERAE_GENUS* | | | -0.39564 | | 0.104123 |
| Aer/Ana | *SICCIRUBRICOCCUS* | | | -0.43209 | | 0.10774 |
| Aer/Ana | *AQUINCOLA* | | | -0.36081 | | 0.108089 |
| Aer/Ana | *UNCLASSIFIED_CYTOPHAGALES_GENUS* | | | 0.360119 | | 0.108811 |
| Aer/Ana | *ROSEOCOCCUS* | | | -0.42753 | | 0.111925 |
| Aer/Ana | *BOSEA* | | | 0.381182 | | 0.118589 |
| Aer/Ana | *ALGORIPHAGUS* | | | -0.47269 | | 0.120699 |
| Aer/Ana | *PARACRAUROCOCCUS* | | | -0.41354 | | 0.125466 |
| Aer/Ana | *DERXIA* | | | -0.40095 | | 0.138566 |
| Aer/Ana | *NOVIHERBASPIRILLUM* | | | -0.44759 | | 0.144545 |
| Aer/Ana | *CHITINOPHAGA* | | | -0.32747 | | 0.147314 |
| Aer/Ana | *PIRELLULA* | | | -0.35561 | | 0.147549 |
| Aer/Ana | *APOSTICHOPUS* | | | -0.70705 | | 0.010128 |
| Aer/Ana | *SKELETONEMA* | | | 0.630377 | | 0.027992 |
| Aer/Ana | *TETRAHYMENA* | | | 0.562994 | | 0.028878 |
| Aer/Ana | *PARAMECIUM* | | | 0.561877 | | 0.029267 |
| Aer/Ana | *TIGRIOPUS* | | | -0.46945 | | 0.031789 |
| Aer/Ana | *PARAMURICEA* | | | -0.46756 | | 0.032577 |
| Aer/Ana | *ICHTHYOPHTHIRIUS* | | | 0.552709 | | 0.032614 |
| Aer/Ana | *PSEUDOCOHNILEMBUS* | | | 0.549228 | | 0.033957 |
| Aer/Ana | *NYMPHAEA* | | | 0.544111 | | 0.036006 |
| Aer/Ana | *LEPEOPHTHEIRUS* | | | -0.46573 | | 0.051427 |
| Aer/Ana | *CRASPEDOSTAUROS* | | | 0.567259 | | 0.054411 |
| Aer/Ana | *ATTHEYA* | | | -0.38152 | | 0.087909 |
| Aer/Ana | *STYLONYCHIA* | | | -0.50671 | | 0.092719 |
| Aer/Ana | *CHAETOCEROS* | | | -0.50033 | | 0.097601 |
| Aer/Ana | *NOCCAEA* | | | 0.440112 | | 0.100646 |
| Aer/Ana | *DITYLUM* | | | -0.36555 | | 0.103194 |
| Aer/Ana | *UNCLASSIFIED_SIPHOVIRIDAE_GENUS* | | | -0.58451 | | 0.022115 |
| Aer/Ana | *CYANOPHAGE* | | | -0.44199 | | 0.044845 |
| Aer/Ana | *UNCLASSIFIED_CAUDOVIRALES_GENUS* | | | -0.42915 | | 0.052221 |
| Aer/Ana | *UNCLASSIFIED_PODOVIRIDAE_GENUS* | | | -0.39876 | | 0.101175 |
| Micropeptin | *UNCLASSIFIED_BATHYARCHAEOTA_GENUS* | | | -0.38504 | | 0.084774 |
| Micropeptin | *UNCLASSIFIED_GEMMATIMONADACEAE_GENUS* | | | 0.625287 | | 0.00552 |
| Micropeptin | *WOLBACHIA* | | | -0.62255 | | 0.005792 |
| Micropeptin | *HERBASPIRILLUM* | | | -0.69002 | | 0.013013 |
| Micropeptin | *GEMMATIMONAS* | | | -0.49778 | | 0.021662 |
| Micropeptin | *ACINETOBACTER* | | | -0.64436 | | 0.023713 |
| Micropeptin | *RHODOFERAX* | | | 0.488081 | | 0.024786 |
| Micropeptin | *PAUCIBACTER* | | | -0.61405 | | 0.033661 |
| Micropeptin | *UNCLASSIFIED_VERRUCOMICROBIACEAE_GENUS* | | | -0.54704 | | 0.03482 |
| Micropeptin | *UNCLASSIFIED_ARMATIMONADETES_GENUS* | | | -0.45981 | | 0.035975 |
| Micropeptin | *ROSEOMONAS* | | | -0.44897 | | 0.04119 |
| Micropeptin | *POLYNUCLEOBACTER* | | | -0.51391 | | 0.050033 |
| Micropeptin | *UNCLASSIFIED_CYTOPHAGALES_GENUS* | | | 0.427287 | | 0.053361 |
| Micropeptin | *PLANKTOPHILA* | | | -0.54341 | | 0.067844 |
| Micropeptin | *UNCLASSIFIED_CHLOROFLEXI_GENUS* | | | -0.40392 | | 0.06938 |
| Micropeptin | *BREVUNDIMONAS* | | | -0.4015 | | 0.071227 |
| Micropeptin | *UNCLASSIFIED_PHYCISPHAERAE_GENUS* | | | -0.43177 | | 0.073576 |
| Micropeptin | *UNCLASSIFIED_PELAGIBACTERALES_GENUS* | | | -0.52941 | | 0.076714 |
| Micropeptin | *NOVIHERBASPIRILLUM* | | | -0.52646 | | 0.078678 |
| Micropeptin | *BELNAPIA* | | | -0.45468 | | 0.088609 |
| Micropeptin | *UNCLASSIFIED_SYNECHOCOCCALES_GENUS* | | | -0.45449 | | 0.088754 |
| Micropeptin | *ROSEOCOCCUS* | | | -0.44649 | | 0.095243 |
| Micropeptin | *UNCLASSIFIED_ACTINOMYCETIA_GENUS* | | | -0.36592 | | 0.102817 |
| Micropeptin | *SICCIRUBRICOCCUS* | | | -0.43354 | | 0.106437 |
| Micropeptin | *UNCLASSIFIED_OPITUTAE_GENUS* | | | -0.42291 | | 0.116282 |
| Micropeptin | *PARACRAUROCOCCUS* | | | -0.41768 | | 0.121342 |
| Micropeptin | *METHYLOBACTERIUM* | | | -0.33637 | | 0.135985 |
| Micropeptin | *SKELETONEMA* | | | 0.741023 | | 0.005825 |
| Micropeptin | *APOSTICHOPUS* | | | -0.68998 | | 0.01302 |
| Micropeptin | *TIGRIOPUS* | | | -0.37636 | | 0.092651 |
| Micropeptin | *PARAMURICEA* | | | -0.36919 | | 0.09955 |
| Micropeptin | *PSEUDO* | | | 0.346343 | | 0.124046 |
| Micropeptin | *CYANOPHAGE* | | | -0.34948 | | 0.120445 |
| Cyclophane-like | *UNCLASSIFIED_BATHYARCHAEOTA_GENUS* | | | -0.56172 | | 0.036579 |
| Cyclophane-like | *WOLBACHIA* | | | -0.76316 | | 0.003881 |
| Cyclophane-like | *OLIGOFLEXUS* | | | 0.827902 | | 0.011155 |
| Cyclophane-like | *UNCLASSIFIED_SYNECHOCOCCALES_GENUS* | | | -0.73805 | | 0.014805 |
| Cyclophane-like | *PAUCIBACTER* | | | -0.80483 | | 0.015971 |
| Cyclophane-like | *UNCLASSIFIED_BDELLOVIBRIONALES_GENUS* | | | 0.710863 | | 0.021193 |
| Cyclophane-like | *UNCLASSIFIED_VERRUCOMICROBIALES_GENUS* | | | -0.6066 | | 0.021449 |
| Cyclophane-like | *UNCLASSIFIED_VERRUCOMICROBIA_GENUS* | | | -0.60468 | | 0.021977 |
| Cyclophane-like | *METHYLOBACTERIUM* | | | -0.59006 | | 0.026336 |
| Cyclophane-like | *BOSEA* | | | 0.628276 | | 0.02868 |
| Cyclophane-like | *UNCLASSIFIED_XANTHOMONADALES_GENUS* | | | 0.754947 | | 0.030359 |
| Cyclophane-like | *UNCLASSIFIED_VERRUCOMICROBIACEAE_GENUS* | | | -0.6773 | | 0.031427 |
| Cyclophane-like | *UNCLASSIFIED_ARMATIMONADETES_GENUS* | | | -0.56951 | | 0.033514 |
| Cyclophane-like | *GEMMATIMONAS* | | | -0.55262 | | 0.040423 |
| Cyclophane-like | *PARABURKHOLDERIA* | | | 0.639625 | | 0.046424 |
| Cyclophane-like | *OHTAEKWANGIA* | | | 0.629832 | | 0.050985 |
| Cyclophane-like | *UNCLASSIFIED_HYPHOMICROBIALES_GENUS* | | | -0.53039 | | 0.05104 |
| Cyclophane-like | *ROSEOMONAS* | | | -0.50999 | | 0.062449 |
| Cyclophane-like | *UNCLASSIFIED_PLANCTOMYCETACEAE_GENUS* | | | 0.50495 | | 0.065528 |
| Cyclophane-like | *UNCLASSIFIED_OPITUTAE_GENUS* | | | -0.57355 | | 0.083004 |
| Cyclophane-like | *UNCLASSIFIED_CHLOROFLEXI_GENUS* | | | -0.46759 | | 0.091806 |
| Cyclophane-like | *SYNECHOCOCCUS* | | | -0.55893 | | 0.093031 |
| Cyclophane-like | *UNCLASSIFIED_ACTINOMYCETIA_GENUS* | | | -0.46576 | | 0.093257 |
| Cyclophane-like | *ACINETOBACTER* | | | -0.62657 | | 0.096445 |
| Cyclophane-like | *RHODOPIRELLULA* | | | 0.624078 | | 0.098181 |
| Cyclophane-like | *POLYNUCLEOBACTER* | | | -0.54784 | | 0.10113 |
| Cyclophane-like | *VAMPIROVIBRIO* | | | 0.61026 | | 0.108112 |
| Cyclophane-like | *UNCLASSIFIED_FLAVOBACTERIACEAE_GENUS* | | | -0.53047 | | 0.114691 |
| Cyclophane-like | *CYANOBIUM* | | | -0.51312 | | 0.129322 |
| Cyclophane-like | *LIMNOHABITANS* | | | -0.42267 | | 0.132157 |
| Cyclophane-like | *CHRYSEOTALEA* | | | 0.447747 | | 0.14439 |
| Cyclophane-like | *TETRAHYMENA* | | | 0.817487 | | 0.00387 |
| Cyclophane-like | *PARAMECIUM* | | | 0.809702 | | 0.004528 |
| Cyclophane-like | *PSEUDOCOHNILEMBUS* | | | 0.803325 | | 0.005124 |
| Cyclophane-like | *ICHTHYOPHTHIRIUS* | | | 0.799409 | | 0.005517 |
| Cyclophane-like | *NYMPHAEA* | | | 0.787085 | | 0.006891 |
| Cyclophane-like | *CRASPEDOSTAUROS* | | | 0.853161 | | 0.007069 |
| Cyclophane-like | *CHAETOCEROS* | | | -0.78724 | | 0.020399 |
| Cyclophane-like | *ATTHEYA* | | | -0.58158 | | 0.029142 |
| Cyclophane-like | *DITYLUM* | | | -0.57856 | | 0.030194 |
| Cyclophane-like | *NOCCAEA* | | | 0.673847 | | 0.032641 |
| Cyclophane-like | *PHAEODACTYLUM* | | | -0.55405 | | 0.0398 |
| Cyclophane-like | *ODONTELLA* | | | -0.5466 | | 0.065931 |
| Cyclophane-like | *GRAMMATOPHORA* | | | -0.45155 | | 0.140597 |
| Cyclophane-like | *PARAMURICEA* | | | -0.41046 | | 0.144905 |
| Cyclophane-like | *FISTULIFERA* | | | -0.4076 | | 0.148004 |
| Cyclophane-like | *TIGRIOPUS* | | | -0.40737 | | 0.148253 |
| Cyclophane-like | *UNCLASSIFIED_SIPHOVIRIDAE_GENUS* | | | -0.5798 | | 0.078941 |
| Cyclophane-like | *UNCLASSIFIED_PODOVIRIDAE_GENUS* | | | -0.525 | | 0.079661 |
| Cyclophane-like | *UNCLASSIFIED_CAUDOVIRALES_GENUS* | | | -0.43758 | | 0.117642 |
| Microviridin B | *UNCLASSIFIED_BATHYARCHAEOTA_GENUS* | | | -0.54366 | | 0.044485 |
| Microviridin B | *UNCLASSIFIED_EURYARCHAEOTA_GENUS* | | | -0.58147 | | 0.130566 |
| Microviridin B | *WOLBACHIA* | | | -0.80994 | | 0.001406 |
| Microviridin B | *UNCLASSIFIED_VERRUCOMICROBIACEAE_GENUS* | | | -0.82399 | | 0.003375 |
| Microviridin B | *PAUCIBACTER* | | | -0.87028 | | 0.00494 |
| Microviridin B | *GEMMATIMONAS* | | | -0.68815 | | 0.006514 |
| Microviridin B | *ACINETOBACTER* | | | -0.84757 | | 0.007873 |
| Microviridin B | *RHODOFERAX* | | | 0.66903 | | 0.008884 |
| Microviridin B | *UNCLASSIFIED_ARMATIMONADETES_GENUS* | | | -0.64982 | | 0.011887 |
| Microviridin B | *UNCLASSIFIED_CHLOROFLEXI_GENUS* | | | -0.63376 | | 0.014948 |
| Microviridin B | *POLYNUCLEOBACTER* | | | -0.71234 | | 0.020805 |
| Microviridin B | *UNCLASSIFIED_SYNECHOCOCCALES_GENUS* | | | -0.70906 | | 0.021674 |
| Microviridin B | *ROSEOMONAS* | | | -0.59088 | | 0.026076 |
| Microviridin B | *HERBASPIRILLUM* | | | -0.75803 | | 0.029302 |
| Microviridin B | *UNCLASSIFIED_ACTINOMYCETIA_GENUS* | | | -0.56311 | | 0.036019 |
| Microviridin B | *UNCLASSIFIED_GEMMATIMONADACEAE_GENUS* | | | 0.601612 | | 0.038501 |
| Microviridin B | *LIMNOHABITANS* | | | -0.54686 | | 0.043 |
| Microviridin B | *BRYOBACTER* | | | -0.64031 | | 0.046116 |
| Microviridin B | *BREVUNDIMONAS* | | | -0.51833 | | 0.057584 |
| Microviridin B | *PLANKTOPHILA* | | | -0.67286 | | 0.067456 |
| Microviridin B | *UNCLASSIFIED_OPITUTAE_GENUS* | | | -0.59786 | | 0.067924 |
| Microviridin B | *SYNECHOCOCCUS* | | | -0.59707 | | 0.06838 |
| Microviridin B | *METHYLOBACTERIUM* | | | -0.48917 | | 0.075866 |
| Microviridin B | *UNCLASSIFIED_VERRUCOMICROBIALES_GENUS* | | | -0.48596 | | 0.078103 |
| Microviridin B | *UNCLASSIFIED_PELAGIBACTERALES_GENUS* | | | -0.64992 | | 0.081073 |
| Microviridin B | *UNCLASSIFIED_VERRUCOMICROBIA_GENUS* | | | -0.47108 | | 0.089076 |
| Microviridin B | *UNCLASSIFIED_FLAVOBACTERIACEAE_GENUS* | | | -0.55257 | | 0.097624 |
| Microviridin B | *RUNELLA* | | | -0.6133 | | 0.105882 |
| Microviridin B | *UNCLASSIFIED_CYTOPHAGALES_GENUS* | | | 0.440241 | | 0.11517 |
| Microviridin B | *BELNAPIA* | | | -0.52926 | | 0.115681 |
| Microviridin B | *NOVIHERBASPIRILLUM* | | | -0.58682 | | 0.126215 |
| Microviridin B | *VAMPIROVIBRIO* | | | -0.57153 | | 0.13887 |
| Microviridin B | *UNCLASSIFIED_RHODOBACTERACEAE_GENUS* | | | -0.4489 | | 0.143238 |
| Microviridin B | *APOSTICHOPUS* | | | -0.97738 | | 2.84E-05 |
| Microviridin B | *SKELETONEMA* | | | 0.831539 | | 0.010493 |
| Microviridin B | *PARAMECIUM* | | | 0.659467 | | 0.038036 |
| Microviridin B | *TETRAHYMENA* | | | 0.656704 | | 0.039138 |
| Microviridin B | *TIGRIOPUS* | | | -0.55241 | | 0.040515 |
| Microviridin B | *ICHTHYOPHTHIRIUS* | | | 0.653081 | | 0.040614 |
| Microviridin B | *NYMPHAEA* | | | 0.651423 | | 0.041302 |
| Microviridin B | *PSEUDOCOHNILEMBUS* | | | 0.641096 | | 0.045763 |
| Microviridin B | *NOCCAEA* | | | 0.588531 | | 0.073478 |
| Microviridin B | *PARAMURICEA* | | | -0.49062 | | 0.07487 |
| Microviridin B | *LEPEOPHTHEIRUS* | | | -0.49739 | | 0.099902 |
| Microviridin B | *EURYTEMORA* | | | 0.442545 | | 0.113059 |
| Microviridin B | *CYANOPHAGE* | | | -0.53873 | | 0.046846 |
| Microviridin B | *UNCLASSIFIED_CAUDOVIRALES_GENUS* | | | -0.48634 | | 0.077838 |
| Microviridin B | *UNCLASSIFIED_SIPHOVIRIDAE_GENUS* | | | -0.54786 | | 0.101118 |
| Microviridin B | *UNCLASSIFIED_PODOVIRIDAE_GENUS* | | | -0.45023 | | 0.141909 |

**Table S4:** Paired environmental variables for the 2014 cyanoHAB

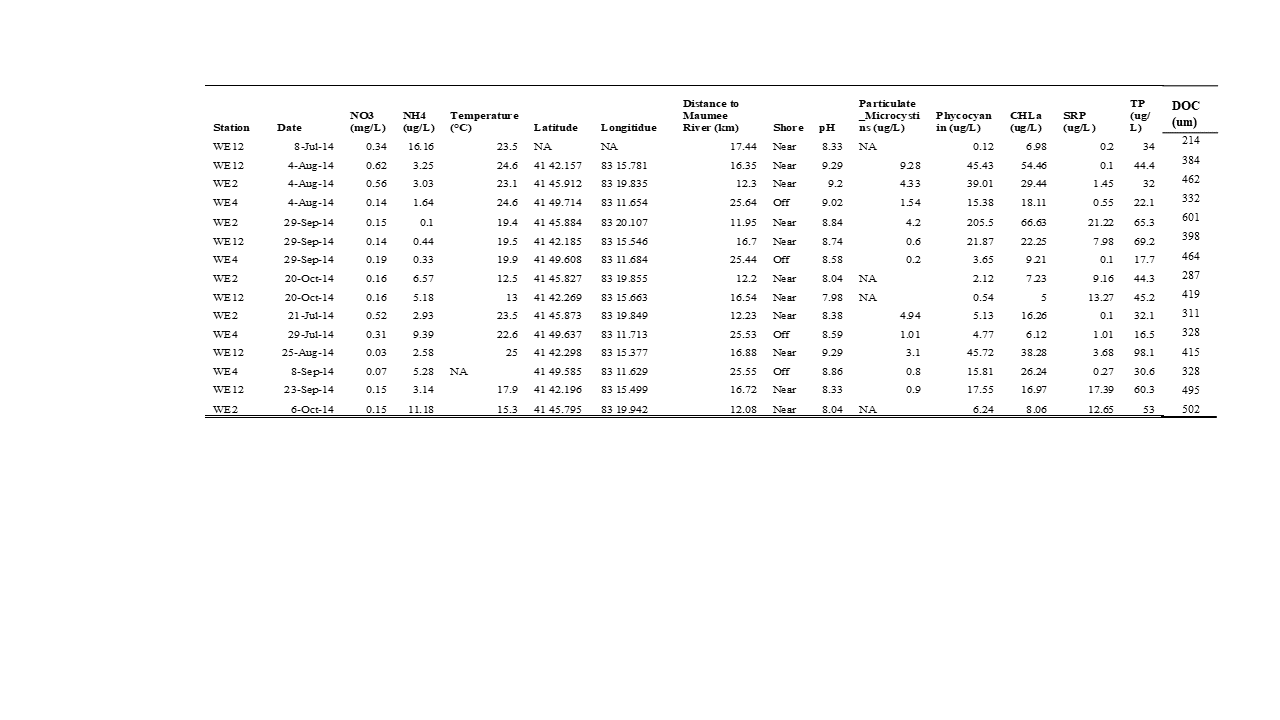
